# Laminar Mechanisms of Saccadic Suppression in Primate Visual Cortex

**DOI:** 10.1101/2021.01.09.426063

**Authors:** Sachira Denagamage, Mitchell P. Morton, John H. Reynolds, Monika P. Jadi, Anirvan S. Nandy

## Abstract

Saccades are a ubiquitous and crucial component of our visual system, allowing for the efficient deployment of the fovea and its accompanying neural resources. Initiation of a saccade is known to cause saccadic suppression, a temporary reduction in visual sensitivity^1, 2^ and visual cortical firing rates^3–6^. While saccadic suppression has been well characterized at the level of perception and single neurons, relatively little is known about the visual cortical networks governing this phenomenon. Here we examine the effects of saccadic suppression on distinct neural subpopulations within visual area V4. We find cortical layer- and cell type-specific differences in the magnitude and timing of peri-saccadic modulation. Neurons in the input layer show changes in firing rate and inter-neuronal correlations prior to saccade onset, indicating that this layer receives information about impending saccades. Putative inhibitory interneurons in the input layer elevate their firing rate during saccades, suggesting they play a role in suppressing the activity of other cortical subpopulations. A computational model of this circuit recapitulates our empirical observations and demonstrates that an input layer-targeting pathway can initiate saccadic suppression by enhancing local inhibitory activity. Collectively, our results provide a mechanistic understanding of how eye movement signaling interacts with cortical circuitry to enforce visual stability.

## INTRODUCTION

As we observe the world around us, our eyes dart from point to point. Each of these shifts in gaze is a saccade – a ballistic movement of both eyes. Saccades significantly improve the efficiency of the primate visual system by allowing us to flexibly deploy our fovea, the dedicated high acuity zone at the center of the retina, towards objects of interest in our environment. However, saccades also pose a substantial challenge for ongoing visual processing, as each saccade produces abrupt and rapid motion across the retina. Therefore, saccades are also accompanied by a temporary reduction in visual sensitivity, termed saccadic suppression, that serves to blunt our perception of this motion^1, 2^. Perhaps reflecting the diminished sensory processing during saccades, the firing rates of neurons in many regions of the visual cortex are also transiently suppressed^3–6^. Yet, despite extensive psychophysical characterization and single neuron electrophysiological analysis, relatively little is known about the circuit level mechanisms underlying saccadic suppression in the visual cortex. One prevailing hypothesis suggests that saccadic suppression is mediated by a corollary discharge signal originating in the brain regions responsible for initiating eye movements^7–9^. Given the synaptic and motor delays accompanying the execution of the saccadic motor command, a coincident corollary discharge to the visual cortex could trigger compensatory mechanisms prior to the start of the eye movement. Consistent with this idea, changes in neural activity before saccade onset have been observed in several visual cortical areas^5^. However, despite clear neural evidence that the visual cortex receives information about upcoming saccades, the pathway by which a saccadic signal arrives in the visual cortex is unknown, and how that signal interacts with local circuitry to mediate saccadic suppression remains unclear.

We examined these questions within visual area V4, a critical hub in the visual processing stream responsible for object recognition^10^. V4 is known to exhibit peri-saccadic suppression in both humans^4^ and macaques^6^, and undergoes dynamic shifts in stimulus sensitivity during the preparation of saccades^11^. To investigate the underlying neural mechanisms, we studied peri- saccadic suppression of neural activity in the context of the laminar cortical circuit, which is composed of six layers with highly stereotyped patterns of intra- and inter-laminar connectivity^12, 13^. In higher-order visual areas, such as V4, layer IV (the *input* layer) is the primary target of projections carrying visual information from lower-order areas, such as V1, V2, and V3^14, 15^. After arriving in the input layer, visual information is then processed by local neural subpopulations as it is sent to layers II/III (the *superficial* layer) and layers V/VI (the *deep* layer), which serve as output nodes in the laminar circuit^12, 16^. In V4, the superficial and deep layers feed information forward to downstream visual areas, such as ST^17^, TEO^18^, and TE^19^. Alongside feedforward input from lower-order visual areas, V4 also receives projections from other cortical regions, such as the prefrontal cortex^15^, as well as from subcortical structures^20^. Accordingly, several pathways targeting V4 could be responsible for relaying information about upcoming saccades: 1) projections from lower visual areas (V1/V2/V3), which predominantly target the input layer^14, 15^, 2) a projection from the frontal eye fields (FEF), which predominantly targets the superficial and deep layers^21, 22^, and 3) a projection from the pulvinar, which predominantly targets the superficial and input layers^23–25^. Neurons in V1^26, 27^, FEF^28–30^, and pulvinar^31–33^ are known to respond to saccades, and the corresponding pathways are therefore strong candidates for initiating suppression in V4.

Given the organization of cortical circuitry, we reasoned that a saccadic signaling pathway should target the site of entry for visual information, the input layer, to suppress visual processing most efficiently. Furthermore, considering the similarities between peri-saccadic stimulus processing and general reductions in visual gain^6, 26^, we considered whether saccadic suppression might be mediated in a manner comparable to visual gain modulation. Specifically, we hypothesized that an elevation in local inhibitory activity, a common mechanism for gain control^34–,36^, functionally suppresses neural processing in V4 during saccades. We sought to confirm our predictions by utilizing a cortical layer-specific recording approach in combination with electrophysiological cell type classification to simultaneously record from six well-defined neural subpopulations in V4.

## RESULTS

### Dissociating the Effects of Vision and Eye Movements

Past approaches for studying peri-saccadic modulation of neural activity have typically involved cued saccade tasks^4, 6, 26, 37^, which risk conflating the neural effects of visual stimulus processing with the neural effects of saccades. To dissociate saccadic signaling from visually induced activity in area V4, we performed a series of spontaneous recordings while two rhesus macaques were seated in the dark. We found that despite their inability to observe specific objects in the environment, both animals continued to make saccades freely under these conditions. Spontaneous neural activity also remained well above the level of noise (Figure S1A). We identified individual saccades by estimating the onset and offset times of ballistic eye movements (see Methods). To remove fixational eye movements (microsaccades), we discarded saccades with amplitudes less than 0.7 degrees of visual angle. Additionally, to prevent the neural effects of one eye movement from overlapping with those of the next, we only included saccades that were separated from one another by at least 500 ms. This approach provided us with a set of 4420 saccades that were used for all subsequent analyses. We evaluated the quality of our eye movement dataset by examining amplitude and velocity traces from individual saccades, as well as by characterizing the main sequence. The main sequence is the approximately linear amplitude-velocity relationship observed across all saccades^38^, which we found held true for our data (Figure 1A).

**Figure 1.**
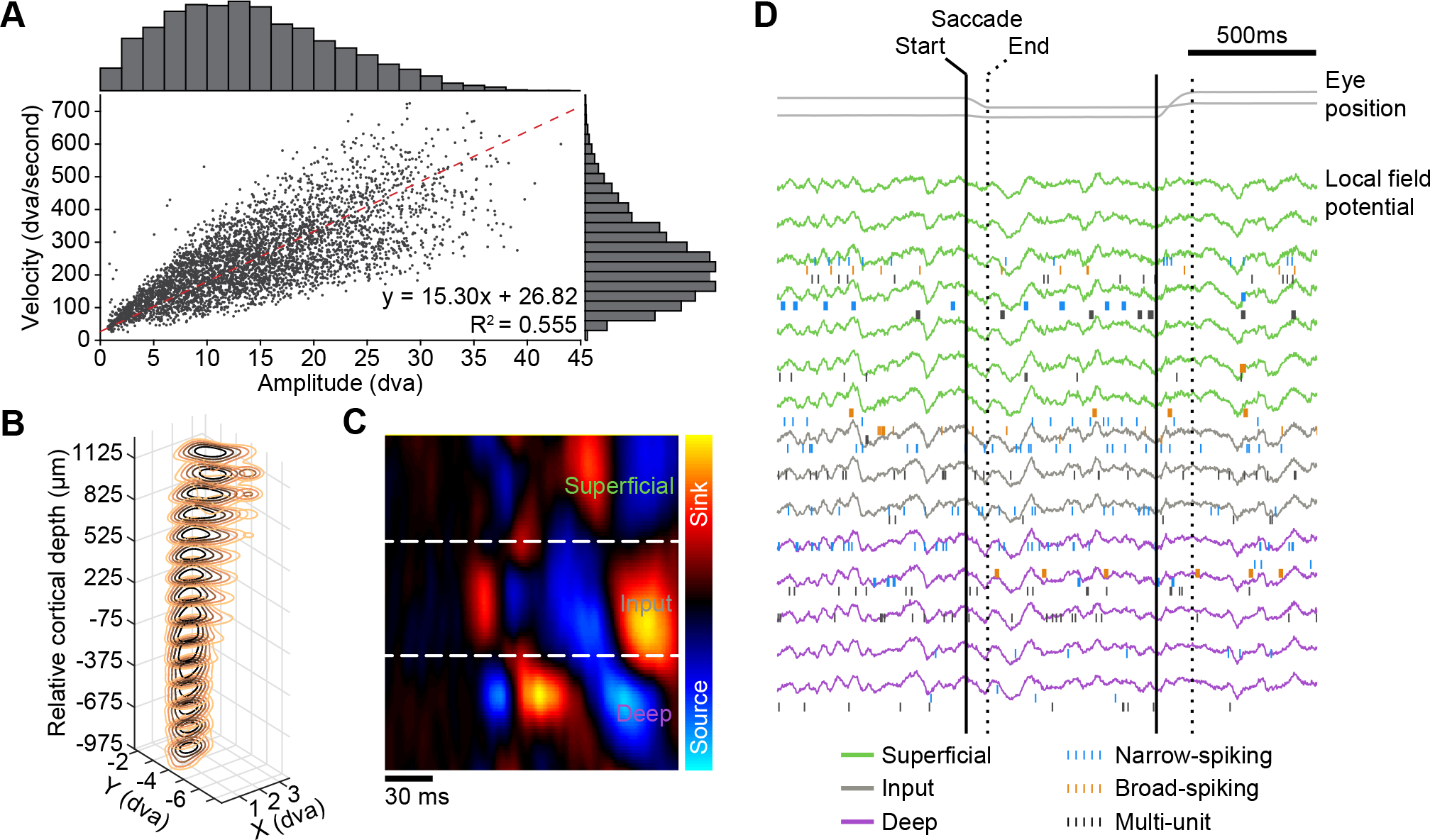
Laminar Recordings and Saccade Identification. (A) The amplitude-velocity relationship (‘main sequence’) for all saccades (*n* = 4420) in the dataset. An approximately linear trend is expected for saccades, as is found here (*R^2^* = 0.555). dva = degrees of visual angle. (B) Stacked contour plot showing spatial receptive fields (RFs) along the laminar probe from an example session. The RFs are well aligned, indicating perpendicular penetration down a cortical column. Zero depth represents the center of the input layer as estimated with current source density (CSD) analysis. (C) CSD displayed as a colored map. The x-axis represents time from stimulus onset; the y-axis represents cortical depth. The CSD map has been spatially smoothed for visualization. (D) Example of continuous eye-tracker and electrophysiological recordings. Eye traces are plotted at the top in grey. LFP traces are plotted by depth below, and spikes are overlaid on the corresponding channel. LFP traces have been color coded by layer identity; spikes have been color coded by cell type.

While the monkeys were making spontaneous saccades, we simultaneously recorded neural activity from well-isolated single-units (*n* = 211), multi-units (*n* = 110), and local field potentials (LFPs) using linear array probes in area V4. The use of recording chambers containing an optically-transparent artificial dura allowed us to make electrode penetrations perpendicular to the cortical surface, and therefore in good alignment with individual cortical columns (see Methods). The alignment of each penetration was confirmed by mapping receptive fields along the depth of cortex, with overlapping receptive fields indicating proper alignment (Figure 1B). Laminar boundaries were identified with current source density (CSD) analysis^39, 40^. The CSD, defined as the second spatial derivative of the LFP, maps the location of current sources and sinks along the depth of the probe. Each layer of the visual cortex produces a characteristic current source-sink pattern in response to visual stimulation, with the input layer producing a current sink followed by a current source, and the superficial and deep layers producing the opposite pattern of activity (Figure 1C). By identifying transitions between current sources and sinks, we assigned laminar identities to each of our recording sites (Figure 1D – LFP traces colored by layer). We also separated well-isolated single-units into two populations based on their waveform duration: narrow-spiking units, corresponding to putative inhibitory interneurons, and broad-spiking units, corresponding to putative excitatory neurons (Figure 1D – spikes colored by unit type)^41–43^. The three cortical layers and two unit types thus gave us six distinct neural subpopulations to consider in subsequent analyses.

### Narrow-Spiking Input Layer Units Show Positive Peri-Saccadic Modulation

To evaluate neural subpopulation-specific responses, we first estimated the peri-saccadic firing rate of well-isolated single-units with a kernel-based approach, in which each spike is convolved with a gaussian kernel^44^ (Figure S1B). We selected the kernel bandwidth on a unit-by-unit basis at each point in time using an optimization algorithm that minimizes mean integrated squared error^45^. When looking at units grouped by cell type, narrow-spiking units exhibited less peri-saccadic suppression than broad-spiking units (Figure 2A). The cause of this difference became clear when the units were further separated by layer. All laminar subpopulations of broad-spiking units displayed peri-saccadic suppression, as did narrow-spiking units in the superficial and deep layers. In contrast, narrow-spiking input layer units elevated their firing rate in response to a saccade (Figure 2B-C; see Figure S2 for additional single-unit examples).

**Figure 2.**
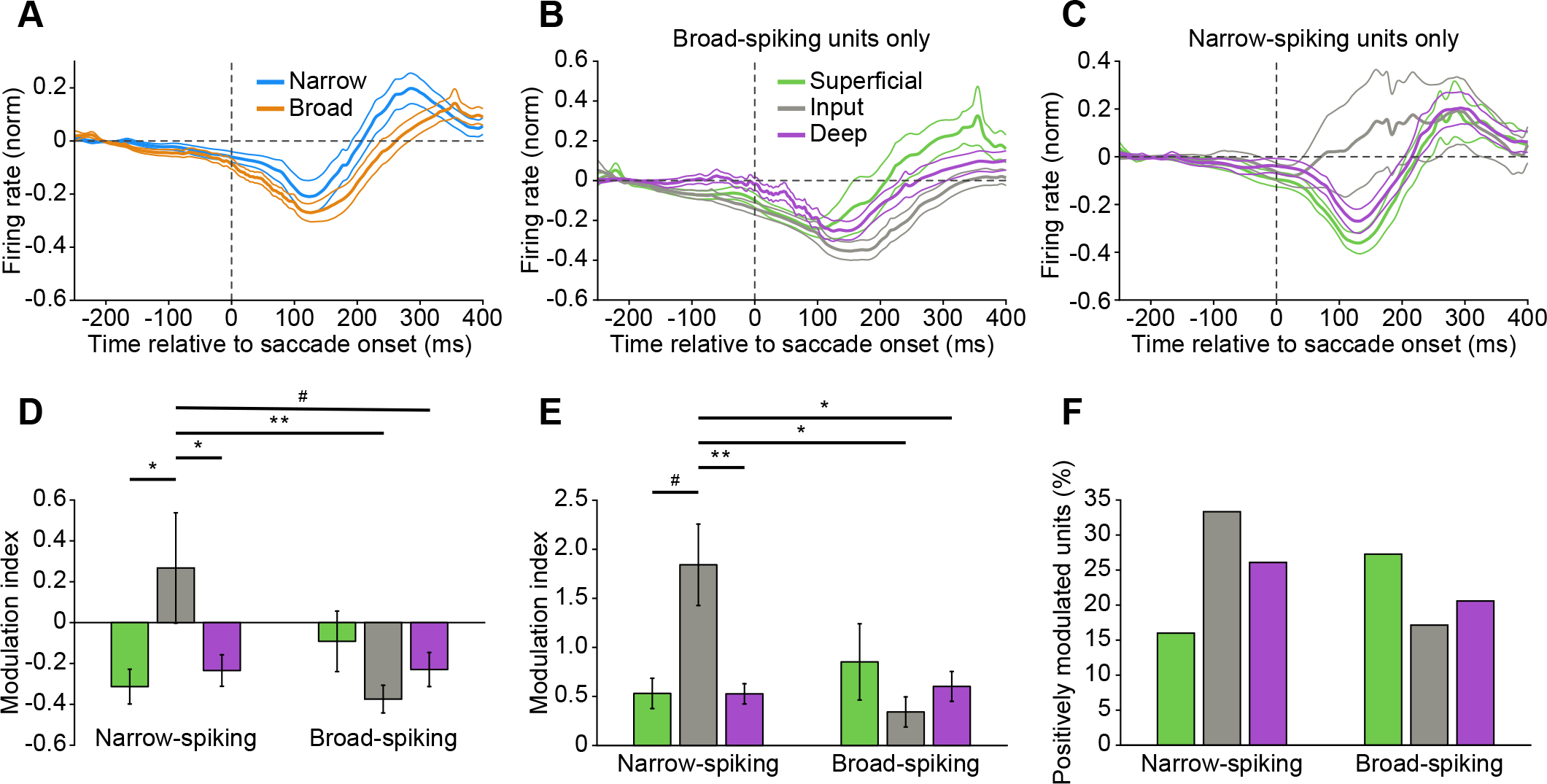
Narrow-Spiking Input Layer Units Display Peri-Saccadic Enhancement of Firing. (A) Average peri-saccadic normalized firing rate for broad- (*n* = 102 units) and narrow-spiking (*n* = 95 units) populations. Thin lines indicate standard error of the mean. (B) As in (A), but including only broad-spiking units from the superficial (*n* = 33), input (*n* = 35), and deep (*n* = 34) layer populations. (C) As in (A), but including only narrow-spiking units from the superficial (*n* = 25 units), input (*n* = 24 units), and deep (*n* = 46 units) layer populations. (D) Average peri-saccadic modulation index for the six neural subpopulations. Two-way ANOVA (*Flayer* = 1.16, *P* = 0.3151; *Fcell type* = 1.84, *P* = 0.1765; *Finteraction* = 5.98, *P* = 0.0030) with Tukey’s multiple comparison tests (narrow-spiking superficial vs narrow-spiking input, *\*P* = 0.0423; narrow-spiking input vs narrow-spiking deep, *\*P* = 0.0495; narrow-spiking input vs broad-spiking input, *\*\*P* = 0.0070; narrow-spiking input vs broad-spiking deep, *^#^P* = 0.0817). Error bars indicate standard error of the mean. (E) Average peri-saccadic modulation index for positively modulated units from the six neural subpopulations. Two-way ANOVA (*Flayer* = 1.87, *P* = 0.1666; *Fcell type* = 2.32, *P* = 0.1355; *Finteraction* = 5.51, *P* = 0.0077) with Tukey’s multiple comparison tests (narrow-spiking superficial vs narrow- spiking input, *^#^P* = 0.0833; narrow-spiking input vs narrow-spiking deep, *\*\*P* = 0.0073; narrow- spiking input vs broad-spiking input, *\*P* = 0.0106; narrow-spiking input vs broad-spiking deep, *\*P* = 0.0384). Error bars indicate standard error of the mean. (F) Proportion of units in each neural subpopulation with a positive peri-saccadic modulation index.

To quantify these observations, we calculated a modulation index that represents the degree to which the firing rate of each unit deviated from its pre-saccadic baseline activity (see Methods). We found that narrow-spiking input layer units were the only group with a positive modulation index (indicating an increase in firing rate), which was significantly different from most other subpopulations (Figure 2D). We also noted that at the single-unit level, each of the six subpopulations had some proportion of positively modulated units. Therefore, the difference in subpopulation level firing rates could be caused by narrow-spiking input layer neurons having a greater average magnitude of positive modulation, or by having a greater proportion of positively modulated units. We determined both to be true. Narrow-spiking input layer neurons displayed a larger magnitude of positive modulation (Figure 2E) and were also more likely to be positively modulated (Figure 2F). We found it particularly striking that, in accordance with our hypothesis, only the narrow-spiking (putative inhibitory interneurons) input layer subpopulation showed an enhancement in firing. This led us to believe that an elevation in input layer inhibitory activity could be responsible for the peri-saccadic suppression of firing within the visual cortex reported previously^3–6^.

### Input Layer Units Respond Prior to Saccades

To gate out visual signals received during a saccade, neurons in the cortical layer initiating suppression would need to be activated prior to the start of the eye movement. To confirm that this was true for the input layer, we performed a timing analysis to determine whether any of the recorded subpopulations show significant changes in firing rate prior to saccade onset. For each single-unit, we found the time at which their activity first deviated outside a 95% confidence interval calculated from their pre-saccadic baseline activity, an event that was marked as the time of first significant response. The results of this approach for six example units (the same example units as in Figure S1B) are illustrated in Figure 3A. While there was response variability at the level of single-units, the input layer subpopulations (both narrow- and broad-spiking) were the only ones whose activity was modulated prior to saccade onset (Figure 3B). This provides strong evidence that the input layer receives information about upcoming saccades with sufficient time to enact changes in activity throughout the cortical column.

**Figure 3.**
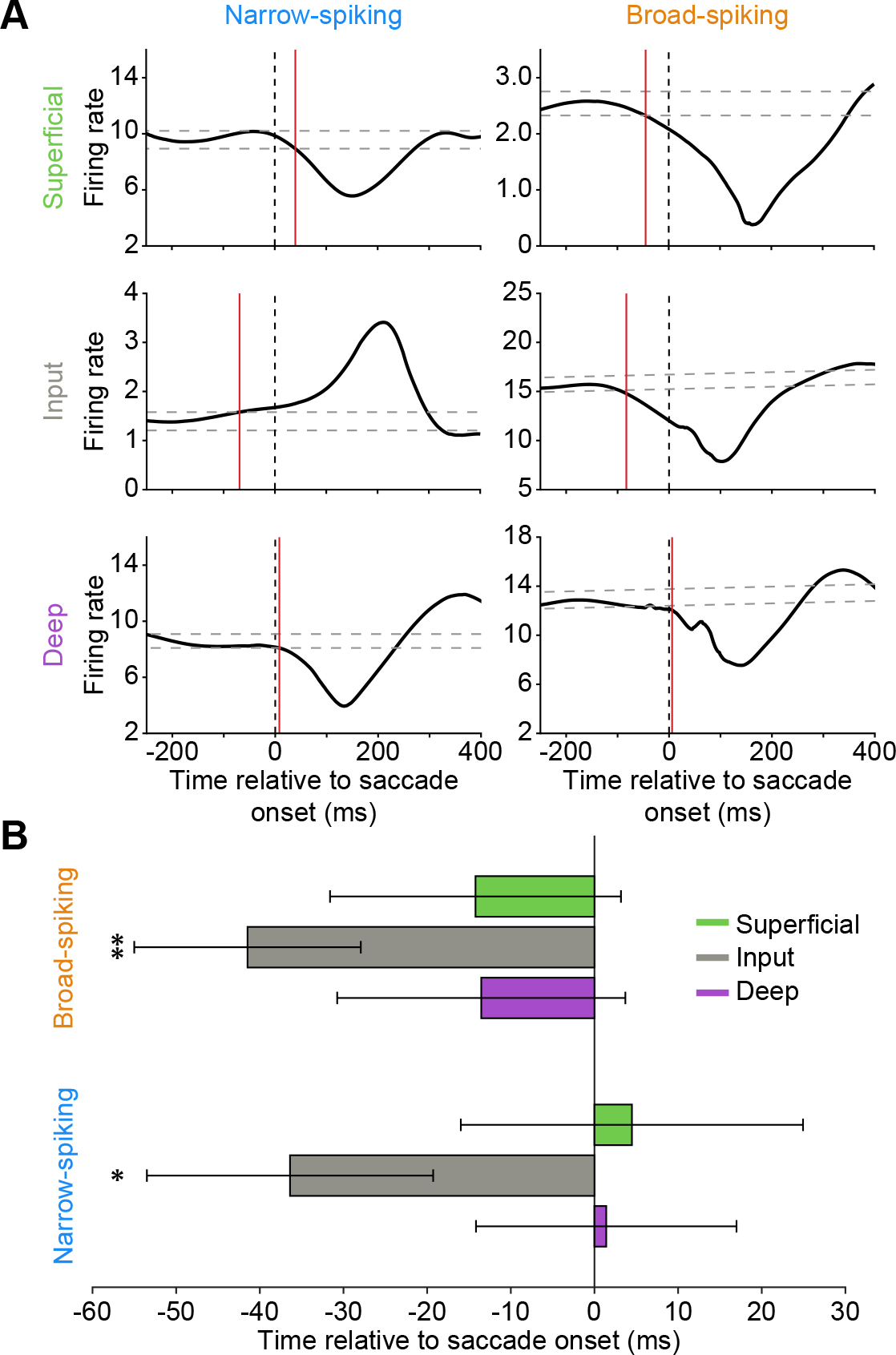
Input Layer Units Display Modulation Prior to Saccade Onset. (A) Six example single-units demonstrating the approach for identifying the time of significant peri-saccadic firing rate modulation. 95% confidence intervals were calculated for the baseline activity of each unit (dashed grey lines) and the first deviation outside of those bounds was marked as the time of first significant response (red line). The example units shown are the same as those from Figure S1B. (B) Average timing of first significant peri-saccadic modulation for the six neural subpopulations. Input layer units, both broad- and narrow-spiking, show modulation prior to saccade onset. Two-tailed one-sample t-test (broad-spiking input layer, *\*\*P* = 0.0049; narrow-spiking input layer, *\*P* = 0.0492). Error bars indicate standard error of the mean.

### Saccades Increase Correlated Variability in the Input Layer

If input layer neurons receive pre-saccadic excitation from a common source, that signaling should also manifest itself as an increase in correlated variability among pairs of input layer units prior to saccade onset. To test this, we computed spike count correlations (SCC) between pairs of simultaneously recorded single- and multi-units as a function of time, and pooled these results into six groups on the basis of their pairwise laminar locations. All pair types showed a substantial increase in SCC around the time of saccades (Figure S3A). We quantified the time at which correlations first began to rise with a bootstrap analysis (see Methods). SCC began increasing prior to saccade onset in input-input pairs (Figure 4A), but after saccade onset in all other pairs (Figure 4B). This pattern of activity is consistent with the pre-saccadic arrival of a signal that is shared among input layer units. The delayed response of all other pair types suggests that they inherit their activity from the input layer. Additionally, the time at which input-input correlations began to rise (∼40 ms before saccade onset) is remarkably similar to the time at which input layer units began to show firing rate modulation (also ∼40 ms before saccade onset), further indicating that these two observations have a common cause: the activation of a saccadic signaling pathway.

**Figure 4.**
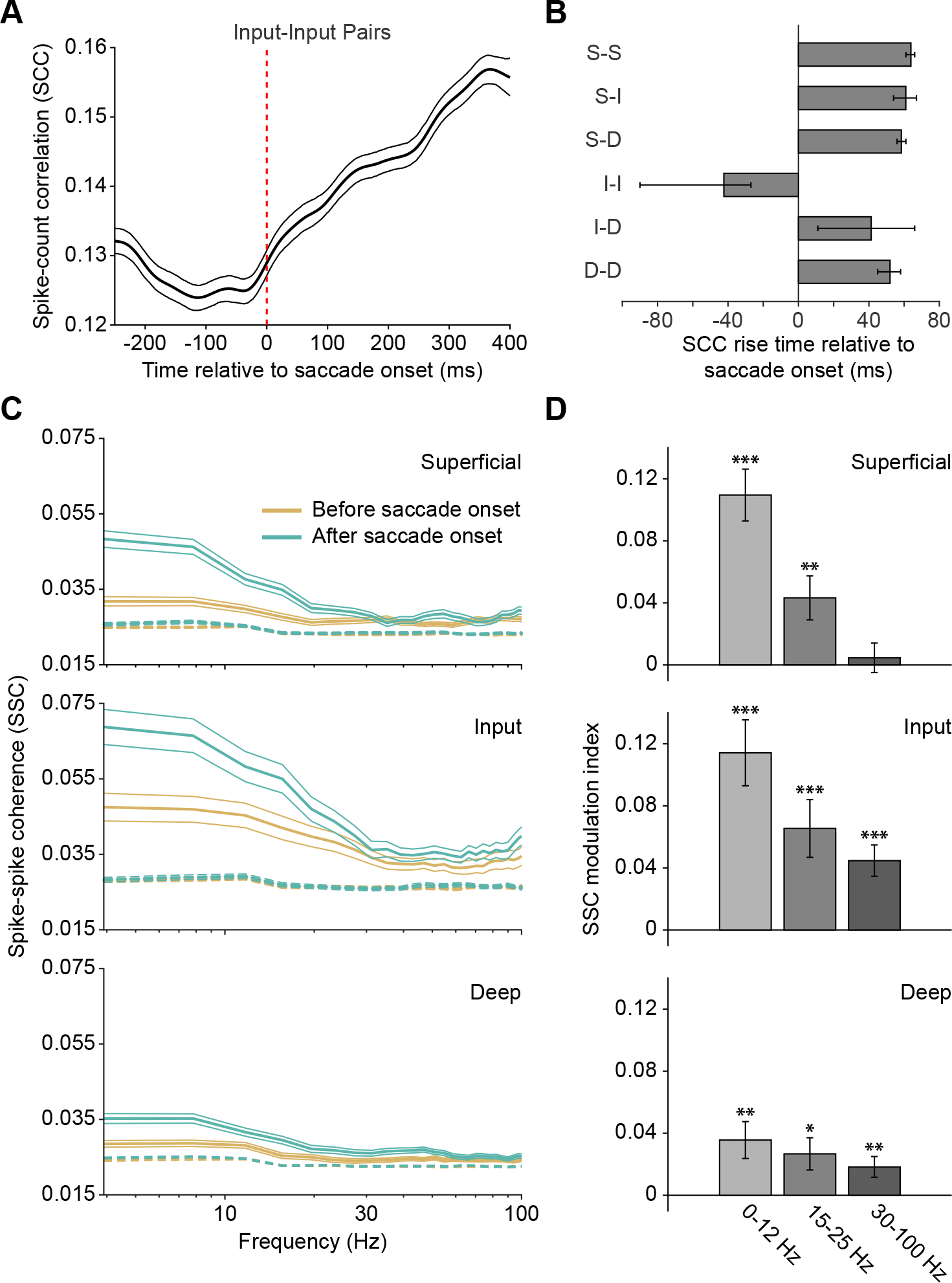
Saccade Onset Increases Correlated Variability Most Prominently in the Input Layer. (A) Spike count correlations as a function of time relative to saccade onset for input-input unit pairs (*n* = 240 pairs). Correlations were calculated with a sliding 201 ms window. Thin lines indicate bootstrapped 95% confidence intervals. (B) Average start time of spike count correlation rise relative to saccade onset. Y-axis labels indicate layer identity of unit pairs (S - superficial; I - input; D - deep). Error bars represent bootstrapped 95% confidence intervals. Input-input pairs begin to rise before saccade onset and before all other pair combinations. All other pairs respond after saccade onset. (C) Average spike-spike coherence between pairs of simultaneously recorded units (single-units and multi-units) before (-200 to 0 ms) and after (0 to 200 ms) saccade onset as a function of frequency. Superficial layer, *n* = 299 pairs; input layer, *n* = 191 pairs; deep layer, *n* = 604 pairs. Thin lines indicate standard error of the mean. Dashed lines indicate spike-spike coherence for shuffled data. (D) Same data as in (C), but averaged across three frequency bands and plotted as a modulation index: (SSCafter - SSCbefore)/(SSCafter + SSCbefore). Two-tailed one-sample t-tests (Superficial 0-12 Hz, *\*\*\*P* = 1.64×10^-10^; Superficial 15-25 Hz, *\*\*P* = 0.0027; Superficial 30-100 Hz, *P* = 0.5468; Input 0-12 Hz, *\*\*\*P* = 2.2131×10^-7^; Input 15-25 Hz, *\*\*\*P* = 5.5458×10^-4^; Input 30-100 Hz, *\*\*\*P* = 1.6503×10^-5^; Deep 0-12 Hz, *\*\*P* = 0.0013; Deep 15-25 Hz, *\*P* = 0.0116; Deep 30-100 Hz, *\*\*P* = 0.0050). Error bars indicate standard error of the mean.

Alongside the expected rise in correlations, we also found a reduction in correlations that began ∼250 ms prior to saccade onset among input-input pairs. Because the timing of this drop precedes saccadic initiation, which begins ∼150 ms before saccade onset^46, 47^, it could not be the result of a corollary discharge signal. Instead, it may reflect the shift in internal state that occurs before saccadic initiation as the subject shifts their attention towards a potential target location^48^. In accordance with this possibility, pupil diameter, a well-established correlate of internal state^49, 50^, begins to ramp up several hundred milliseconds prior to saccade onset (Figure S3B)^51^. Similar shifts in internal state are known to be accompanied by comparable reductions in correlated variability^50, 52–54^.

To further characterize changes in coordinated peri-saccadic activity, we examined differences in spike-spike coherence (SSC) before and after saccade onset. Spike-spike coherence is a measure of the degree to which the activity of two signals, in this case a pair of simultaneously recorded single- and/or multi-units, fluctuates together across a range of frequencies. We found that saccade onset increases wide-band coherence most prominently in the input layer, with weaker, but still significant, effects in the superficial and deep layers (Figure 4C). When quantified as a modulation index (Figure 4D), we observed significant modulation in all layers at all frequencies, with the exception of the 30-100 Hz band in the superficial layer. The input layer exhibits larger increases in coherence, particularly at higher frequencies, while the deep layer experiences smaller increases, particularly at lower frequencies. Collectively, these data suggest that saccade onset leads to a temporary elevation of correlated activity, which manifests itself most prominently in the input layer. The timing and magnitude of changes in correlated variability, measured with both SCC and SSC, further suggests that an external signal arrives prior to saccade onset within the input layer before then propagating to the other layers.

### Saccades Increase the Strength of Low-Frequency Wide-Band Activity

Increases in coordinated activity at the level of single neurons can result in an elevation of population-level wide-band activity. To determine whether this held true for saccades, we calculated the spectral power, a frequency-resolved measure of signal strength, of the LFP around the time of saccade onset. Since LFP signals are highly correlated across the depth of the cortex, we chose to perform this analysis by collapsing across laminar compartments. Saccade onset was associated with a large elevation in low frequency power, and a slight reduction in high frequency power (Figure 5A). Comparing the power spectrum before and after saccade onset, we found that these differences were significant in all three frequency bands tested, with an elevation in the 0-12 Hz and 15-25 Hz bands and a reduction in the 30-80 Hz band after saccade onset (Figure 5B). Next, we sought to investigate whether the increased strength of low-frequency wide-band activity entrains spiking units, as reported previously in other contexts^55–57^. We found that saccade onset significantly increases low-frequency spike-LFP phase locking (Figure 5C), as measured by pairwise phase consistency (PPC). These results indicate that saccade onset increases the strength of low-frequency population-level activity, which could in turn entrain the activity of local neurons.

**Figure 5.**
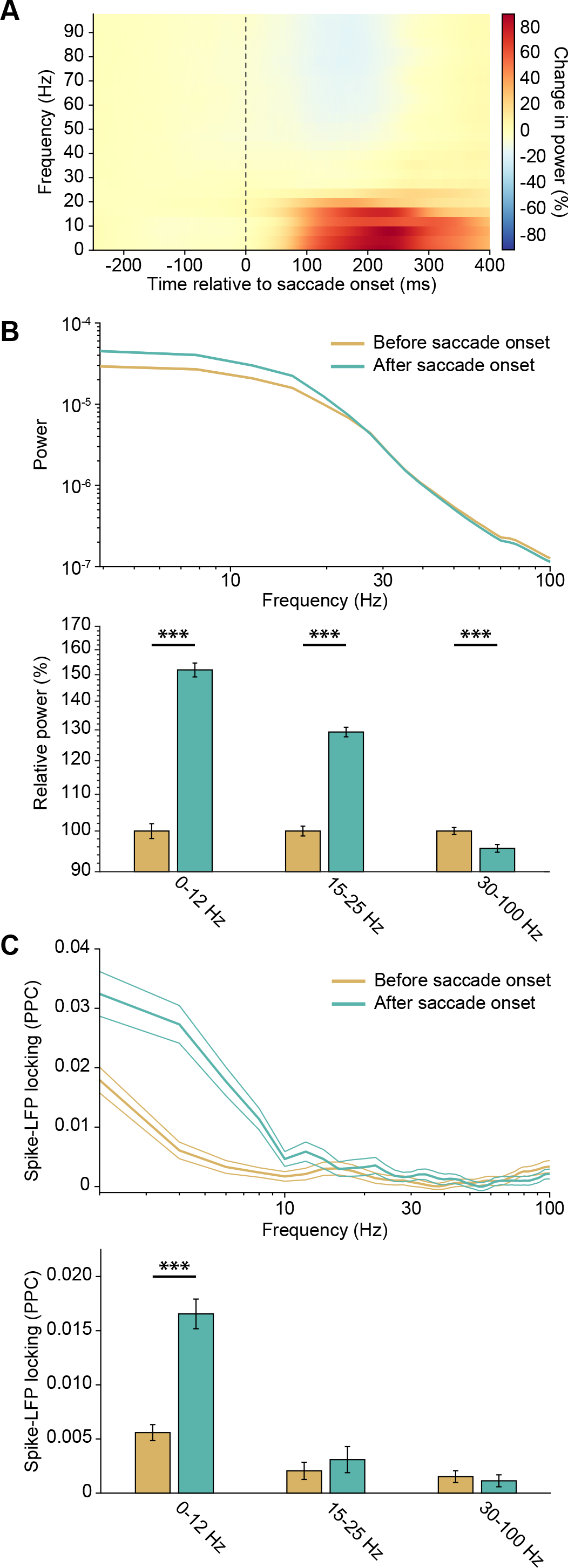
Saccade Onset Increases Low Frequency Power and Spike-LFP Locking. (A) Time-frequency representation (spectrogram) of LFP power around saccade onset. Signal strength is represented as percent change in power (i.e. the raw power at each timepoint is divided by the average power in the baseline period, defined here as 250 to 150 ms before saccade onset). (B) Top: Average power spectra before (-200 to 0 ms) and after (0 to 200 ms) saccade onset. Thin lines indicate standard error of the mean. Bottom: Same data as in top, but averaged across three frequency bands and normalized to data before saccade onset for visualization. Two tailed paired- sample t-tests (0-12 Hz, *\*\*\*P* = 3.4669×10^-92^; 15-25 Hz, *\*\*\*P* = 9.3690×10^-74^; 30-100 Hz, *\*\*\*P* = 2.7322×10^-9^). Error bars indicate standard error of the mean. (C) Top: Average spike-LFP locking spectra before (-200 to 0 ms) and after (0 to 200 ms) saccade onset. Thin lines indicate standard error of the mean. Bottom: Same data as in top, but averaged across three frequency bands. Two tailed paired-sample t-tests (0-12 Hz, *\*\*\*P* = 1.7147×10^-19^; 15- 25 Hz, *P* = 0.1598; 30-100 Hz, *P* = 0.1091). Error bars indicate standard error of the mean.

### A Computational Model Demonstrates that Activation of an Input Layer-Targeting Projection is Sufficient for Inducing Saccadic Suppression

Given that input layer units exhibit significant changes in firing properties prior to saccade onset, we hypothesized that a projection targeting this layer could serve as the initiator of saccadic suppression within area V4. To determine whether such a projection would be sufficient for initiating suppression across all cortical layers, as well as to gain further insight into the associated circuit-level mechanisms, we developed a computational model of peri-saccadic activity changes within a cortical column. Our model draws inspiration from previous cortical circuit models that describe population-specific firing rate changes in response to the activation of an external input^58^. To simulate cortical circuit-level dynamics more faithfully, our model consists of six interconnected populations across three layers of the cortex (Figure 6A, left). Each cortical layer contains an excitatory (E) and inhibitory (I) population, which project to each other as well as to themselves. Given evidence of multiple operating regimes for a local E-I network in the visual cortex, connections within a layer were tuned to allow the local E-I network to switch in and out of an inhibition-stabilized network (ISN) regime^59, 60^: the baseline regime of the network is non-ISN, but direct external input to the E population switches it to ISN (see Methods; Figure S4A-C). Connections between layers exist in the form of projections from E neurons in the source layer to both E and I neurons in the target layer, with E-E and E-I connections tuned to affect a net increase in E activity in the target layer in response to an increase in E activity in the source layer. In our model, the input layer projects to the superficial layer, which then projects to the deep layer; this connectivity motif highlights the primary pathway for information flow within a visual cortical column^13, 61^. Considering our empirical firing rate analysis, which found that both broad- and narrow-spiking units in the input layer show responses prior to saccade onset (Figure 3B), we chose to model the input layer as the primary recipient of the ‘saccadic signal’ input. We tuned the strength of the excitatory ‘saccadic signal’ pathway to the local E and I populations in this layer such that its activation affected a net suppression of the E activity.

**Figure 6.**
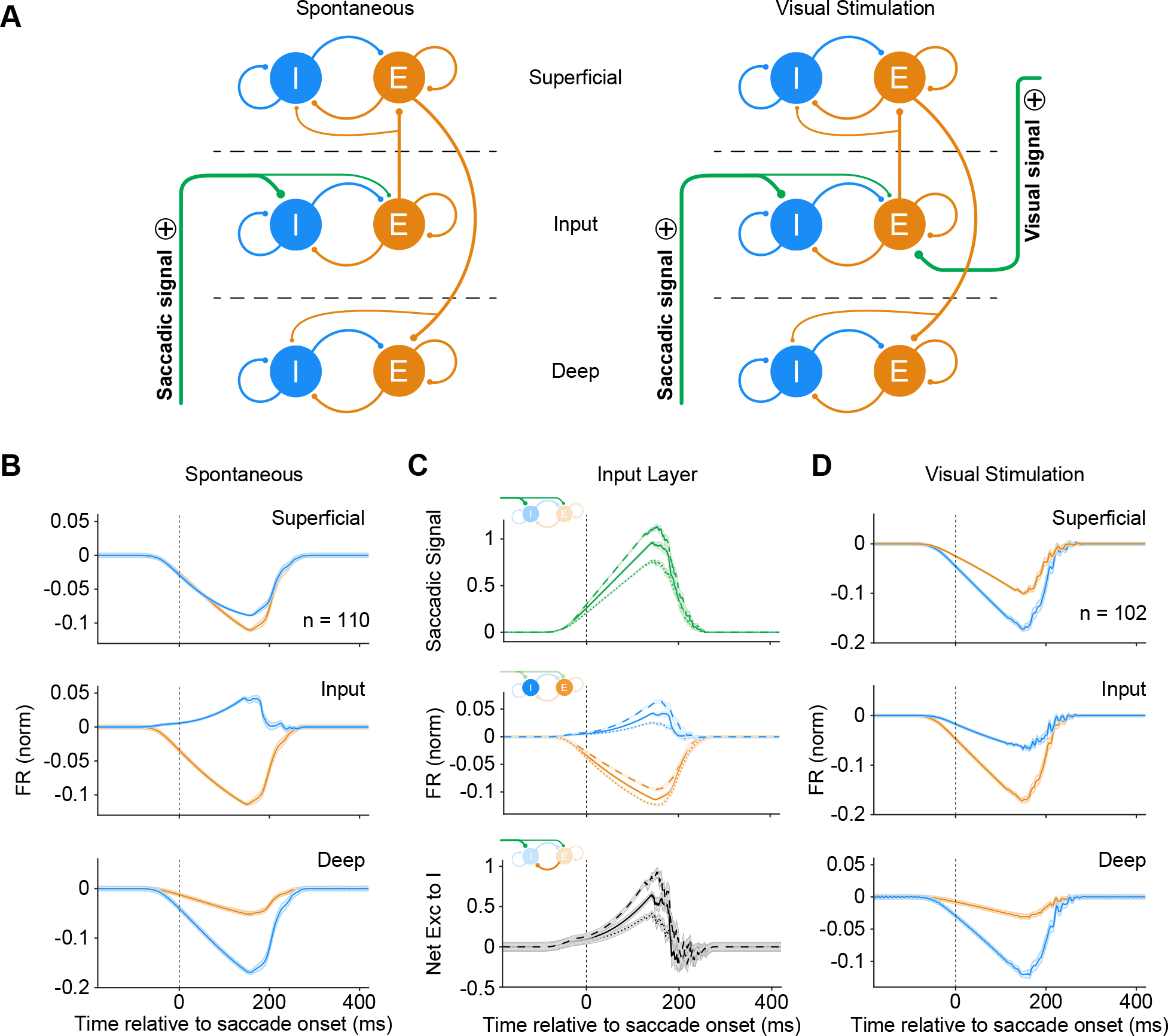
A Computational Model Confirms that Activation of an Input Layer-Targeting Projection Induces Saccadic Suppression. (A) Connectivity between populations in our computational model. Excitatory (E) and inhibitory (I) populations within each layer project to each other as well as themselves. The input layer excitatory population projects to the superficial layer, while the superficial excitatory population projects to the deep layer. In our model of spontaneous saccades (left), we activate an external excitatory projection that selectively targets the input layer. In our model of saccades executed in the presence of visual stimuli (right), we add an additional input that selectively targets input layer excitatory neurons. (B) Normalized peri-saccadic firing rate of simulated neural subpopulations during spontaneously executed saccades (*n* = 110 simulated saccades). The input layer inhibitory subpopulation shows enhancement of firing, while all other subpopulations show suppression. (C) Further dissection of input layer activity. (Top) The saccadic signal arriving in the input layer was simulated with a ramping function. Represented here are saccadic signals of three different strengths. From weakest to strongest, these are represented by dotted, continuous, and dashed lines, both here and in the following subpanels. (Middle) The excitatory and inhibitory input layer subpopulation firing rates in response to three saccadic signals of varying strength. (Bottom) Net excitatory drive onto the inhibitory input layer subpopulation in response to three saccadic signals of varying strength. The excitatory drive is the sum of drive from the external saccadic input and drive from the local excitatory population. (D) Normalized peri-saccadic firing rate of simulated neural subpopulations during saccades executed in the presence of visual stimulation (*n* = 102 simulated saccades). Visual stimulation is represented by the activation of a second input, which selectively targets the input layer excitatory subpopulation. Here, all subpopulations within the cortical column show peri-saccadic suppression.

In response to the activation of a saccadic signal of sufficient strength slightly before the time of saccade onset, the E input layer population rate was suppressed. In addition, our model simultaneously recapitulated our experimental findings in the input layer I population (enhancement), as well as the E/I populations in other layers (suppression) by virtue of the inter- and intra-layer connectivity (Figure 6B, S4B). Since local E activity is a major source of excitation to the I population within a layer, we next explored the role of the saccadic signal in sustaining input layer I activity despite a concurrent decline in local E activity. We explored three scenarios with saccadic signals of varying magnitudes ramping up and down over a fixed period (Figure 6C, top). We found that the increase in I activity in the input layer indeed depended on the strength of the saccadic signal and was sustained at sufficiently high magnitudes. Examination of the contribution of different excitatory pathways to the I population showed that sufficiently high saccadic signal magnitude led to an increase in net excitation onto the I population that exceeded the loss in excitation resulting from the reduction of local E firing rates (Figure 6C, middle and bottom). Our model thus demonstrates that an input layer-specific saccadic signal of adequate strength can increase local I activity, which is sufficient for inducing suppression across the depth of the cortex. These findings correspond closely to our empirical observations. The model further predicted that the time course of the increase in input layer I activity was dependent on the baseline operating regime of the E-I network. When the network was robustly within the non-ISN regime, I activity increased immediately following saccadic signal onset. However, when the network was close to the switching point between the two regimes, the I activity showed little change below a saccadic signal of sufficient magnitude, even when the local E activity showed significant suppression (Figure S4E). This latter scenario corresponds closely to our empirical observations in the input layer (Figure 2B-C), suggesting that the baseline state of the cortical network lies close to the boundary between regimes.

Our experimental paradigm examined peri-saccadic neural activity in the absence of visual input. While this allowed us to investigate the cortical dynamics that are purely a result of the saccadic signal and avoid potential stimulus-induced artifacts in our recordings, it leaves open the question of how activity patterns may change in the presence of visual stimulation. To explore this further, we added a second external excitatory input (‘visual signal’) to the input layer E population (Figure 6A, right), which is the major target of feedforward visual input into V4^14^. The addition of this second input shifted the local E-I network into an ISN regime^59^, in which E and I activity increase or decrease concurrently. As a result, simultaneous activation of the visual signal led to a net reduction in the firing rate of both E and I input layer populations (Figure 6D, S4C). This produced a similar reduction in excitatory drive to the superficial and deep layers, where firing was again suppressed. These results are consistent with prior reports of suppression from uncategorized neurons in the visual cortex in the presence of visual stimulation^3, 5, 6^.

## DISCUSSION

To contextualize our findings regarding neurophysiological saccadic suppression in area V4, we must also consider how this phenomenon may relate to perceptual saccadic suppression, which has been extensively studied. Perceptual suppression has disputed origins: some groups have argued for extra-retinal origins^62^ such as corollary discharge, while others have argued for visual masking mechanisms^63, 64^. There is also significant uncertainty about the prevalence of suppression in the dorsal (motion-detection) versus ventral (object-recognition) visual streams, with cortical suppression having been studied predominantly in the dorsal stream^3, 5^ despite conflicting psychophysical evidence on the impact of saccades on motion perception^63, 65^. Recent studies have demonstrated that saccadic suppression effects are prevalent in the ventral stream as well^6, 26^. Given this body of work, it is important to note that the mechanism described here is not likely to be solely responsible for neural suppression across all visual areas, nor is it likely to be the singular initiator of perceptual suppression during saccades. Instead, perceptual suppression almost certainly arises from a coordinated wave of effects across visual and non-visual areas around the time of saccades. This may also explain why neural suppression observed in other studies^6, 26^, as well as our own, often outlasts perceptual suppression – while some visual areas are suppressed, other brain regions may actually act to shorten perceptual suppression instead of initiating or lengthening it.

The results described here provide evidence that the input layer of area V4 receives the earliest information about upcoming saccades. Units in the input layer showed changes in firing rate prior to saccade onset, while units in other layers did not. In addition, elevated spike-count correlations, a common byproduct of activating a shared input to a neural population^66–69^, were observed earliest among pairs of input layer neurons. Consistent with a single input source (i.e. projection) producing both phenomena, changes in the firing rate and correlational structure of input layer units were initiated with similar timing, approximately 40 ms before saccade onset. These convergent results indicate that the saccadic signaling pathway in question predominantly targets the input layer, which thereby allows us to infer its source. Given the lack of pre-saccadic modulation in the superficial and deep layers, it is unlikely that a projection from the FEF is responsible for the observed suppression. Likewise, it is doubtful that suppression is fed forward by projections from lower visual areas (V1/V2/V3), as such a mechanism would not explain the enhancement of narrow-spiking input layer firing rates in V4. Furthermore, to our knowledge, there have been no clear reports of saccadic suppression in V2 or V3, and while V1 is known to be suppressed during saccades, it does not exhibit pre-saccadic modulation^26^, and could therefore not be responsible for the changes in activity described here. Thus, we propose that saccadic suppression in V4 is likely initiated via the pulvinar, which sends projections to the input layer of _V4_^23-25^.

Several lines of evidence are consistent with this view. First, the pulvinar is known to be the target of highly organized projections from the superior colliculus^70^, which is a critical component of the saccadic initiation network^71, 72^. Second, peri-saccadic modulation has been observed in the pulvinar of macaques, where neurons respond to saccades made in the dark without visual stimulation^32, 33^. Neurons in the pulvinar of cats are able to distinguish between internally generated saccades and simulated saccades produced with image motion, demonstrating that they encode the underlying motor command^31^. Lastly, the pulvinar is known to play a crucial role in spatial attention^73–75^, which is necessary for the proper execution of saccades towards target stimuli; indeed, temporary inactivation of the pulvinar produces significant deficits in saccadic target selection and execution^76, 77^. Thus, pulvinar neurons encode the necessary information about saccadic motor planning to be able to provide a meaningful pre-saccadic signal to their efferent targets in the visual cortex.

We also found that narrow-spiking units in the input layer showed a peri-saccadic enhancement in firing. Narrow-spiking units are thought to correspond to parvalbumin-expressing (PV) inhibitory interneurons^78, 79^, and input layer PV interneurons are known to receive strong and selective thalamocortical excitation.^80–82^ This led us to speculate that an elevated level of inhibition within the input layer could suppress local excitatory neurons, as well as neurons in other cortical layers (Figure 6A). To explore this hypothesis, we developed a six-population firing rate model of the visual cortex and its modulation during saccades. The model demonstrated that pre-saccadic activation of an input layer-targeting projection is sufficient for suppressing signaling within a cortical column. A compelling aspect of our model is its simplicity – with straightforward inter-population connectivity rules, we were able to both replicate our empirical findings as well as reconcile the opposing patterns of activity observed in the input layer subpopulations. The model also reproduces trends in our experimental observations on the relative time course of broad and narrow spiking activity under a set of conditions wherein the input layer operates on the cusp of ISN and non-ISN operating regimes. Given these results, we propose that inhibitory input layer neurons initiate saccadic suppression and gate the processing of visual information entering the local cortical network.

We also observed increases in correlated activity across all cortical layers during saccades, in the form of elevated spike-count correlations, spike-spike coherence, low-frequency power, and low frequency spike-phase locking. Comparable increases in neural correlation have been associated with diminished stimulus sensitivity in multiple model organisms across a variety of sensory modalities^50, 83–86^. This may be a result of heightened correlations functionally limiting the ability of a neural population to encode information, as suggested by previous experimental and theoretical work^87–90^. Similarly, saccadic suppression at the level of neurons and circuits is also accompanied by a behaviorally significant reduction in stimulus detection capabilities^1, 2^, indicating that the increases in correlated variability reported here may reflect or contribute to that deficit.

To isolate the effects of the corollary discharge signal in initiating saccadic suppression in V4, we chose to examine neurophysiological responses to saccades executed in the dark, a paradigm that has been employed extensively by previous studies^6, 31, 91^. Under these conditions, the time course of suppression that we observed was similar to reports of V4 activity during saccades made with visual stimulation^6^, suggesting that the saccadic signaling network remains intact in dark conditions. Our model demonstrates that a network regime change, which has been shown to occur when the cortical network is actively engaged in sensory processing^59, 60^, can flip the modulation of the inhibitory input layer subpopulation in the presence of visual stimulation. This result is consistent with the lack of evidence for subpopulation-specific differences in modulation from previous studies employing cued saccade tasks. In summary, our study provides the first mechanistic understanding of peri-saccadic neural dynamics in a defined cortical circuit.

## ACKNOWLEDGEMENTS

This research was supported by NIH R01 EY021827 to JHR and ASN, NARSAD Young Investigator Grant, Ziegler Foundation Grant and Yale Orthwein Scholar Funds to ASN, NIH R00 EY025026 to MPJ, and by NEI core grants for vision research P30 EY019005 to the Salk Institute and P30 EY026878 to Yale University. We would like to thank Catherine Williams and Mat LeBlanc for excellent animal care.

## AUTHOR CONTRIBUTIONS

SD & ASN conceptualized the project. ASN collected the data and supervised the project. SD analyzed the data, with assistance from MPM and ASN. MPJ developed the computational model. SD, ASN, MPJ, and JHR wrote the manuscript.

## METHODS

### Surgical Procedures

Surgical procedures have been described in detail previously^92, 93^. In brief, an MRI-compatible, low-profile titanium chamber was placed over the pre-lunate gyrus on the basis of preoperative MRI imaging in two rhesus macaques (right hemisphere in monkey A, left hemisphere in monkey C). The native dura mater was then removed, and a silicone-based, optically clear artificial dura (AD) was inserted, resulting in an optical window over dorsal V4. All procedures were approved by the Institutional Animal Care and Use Committee and conformed to NIH guidelines.

### Electrophysiology

At the beginning of each recording session, a plastic insert with an opening for targeting electrodes was lowered into the chamber and secured. This served to stabilize the recording site against cardiac pulsations. Neurons were recorded from cortical columns in dorsal V4 using 16- channel linear array electrodes (‘‘laminar probes’’; Plexon, Plexon V-probe). The laminar probes were mounted on adjustable x-y stages attached to the recording chamber and positioned over the center of the pre-lunate gyrus under visual guidance through a microscope (Zeiss). This ensured that the probes were maximally perpendicular to the surface of the cortex and thus had the best possible trajectory to make a normal penetration down a cortical column. Across recording sessions, the probes were positioned over different sites along the center of the gyrus in the parafoveal region of V4 with receptive field eccentricities between 2 and 7 degrees of visual angle. Care was taken to target cortical sites with no surface micro-vasculature and, in fact, the surface micro-vasculature was used as reference so that the same cortical site was not targeted across recording sessions. The probes were advanced using a hydraulic microdrive (Narishige) to first penetrate the AD and then through the cortex under microscopic visual guidance. Probes were advanced until the point that the top-most electrode (toward the pial surface) registered LFP signals. At this point, the probe was retracted by about 100–200 mm to ease the dimpling of the cortex due to the penetration. This procedure greatly increased the stability of the recordings and also increased the neuronal yield in the superficial electrodes.

The distance from the tip of the probes to the first electrode contact was either 300 mm or 700 mm. The inter-electrode distance was 150 mm, thus negating the possibility of recording the same neural spikes in adjacent recording channels. Neuronal signals were recorded extra- cellularly, filtered, and stored using the Multi-channel Acquisition Processor system (Plexon). Neuronal signals were classified as either multi-unit clusters or isolated single-units using the Plexon Offline Sorter program. Single-units were identified based on two criteria: (1) if they formed an identifiable cluster, separate from noise and other units, when projected into the principal components of waveforms recorded on that electrode and (2) if the inter-spike interval (ISI) distribution had a well-defined refractory period. Single-units were classified as either narrow-spiking (putative interneurons) or broad-spiking (putative pyramidal cells) based on methods described in detail previously^41^. Specifically, only units with waveforms having a clearly defined peak preceded by a trough were potential candidates. Units with trough-to-peak duration less than 225 ms were classified as narrow-spiking units; units with trough- to-peak duration greater than 225 ms were classified as broad-spiking units.

Data were collected over 17 sessions (8 sessions in monkey A and 9 in monkey C), yielding a total of 211 single-units (99 narrow-spiking, 112 broad-spiking) and 110 multi-unit clusters.

### Recording

Stimuli were presented on a computer monitor placed 57 cm from the eye. Eye position was continuously monitored with an infrared eye tracking system (ISCAN ETL-200). Trials were aborted if eye position deviated more than 1 degree of visual angle from fixation. Experimental control was handled by NIMH Cortex software (http://www.cortex.salk.edu/).

### Receptive Field Mapping

At the beginning of each recording session, neuronal RFs were mapped using subspace reverse correlation in which Gabor (eight orientations, 80% luminance contrast, spatial frequency 1.2 cycles/degree, Gaussian half-width 2 degrees) or ring (80% luminance contrast) stimuli appeared at 60 Hz while monkeys fixated. Each stimulus appeared at a random location selected from an 11 x 11 grid with 1 degree spacing in the appropriate visual quadrant. Spatial receptive maps were obtained by applying reverse correlation to the evoked LFP signal at each recording site. For each spatial location in the 11 x 11 grid, we calculated the time-averaged power in the stimulus-evoked LFP (0–200 ms after each stimulus flash) at each recording site. The resulting spatial map of LFP power was taken as the spatial RF at the recording site. For the purpose of visualization, the spatial RF maps were smoothed using spline interpolation and displayed as stacked contours plots of the smoothed maps (Figure 1B). All RFs were in the lower visual quadrant (lower left in monkey A and lower right in monkey C) and with eccentricities between 2 and 7 degrees of visual angle.

### Current Source Density Mapping

In order to estimate the laminar identity of each recording channel, we used a CSD mapping procedure^39^. Monkeys maintained fixation while 100% luminance contrast ring stimuli were flashed (30 ms), centered at the estimated RF overlap region across all channels. The size of the ring was scaled to about three-quarters of the estimated diameter of the RF. CSD was calculated as the second spatial derivative of the flash-triggered LFPs. The resulting time-varying traces of current across the cortical layers can be visualized as CSD maps (Figure 1C). Red regions depict current sinks in the corresponding region of the cortical laminae; blue regions depict current sources. The input layer (Layer 4) was identified as the first current sink followed by a reversal to current source. The superficial (Layers 1–3) and deep (Layers 5 and 6) layers had opposite sink-source patterns. LFPs and spikes from the corresponding recording channels were then assigned to one of three layers: superficial, input, or deep.

### Spontaneous Recordings

In the main experiment, all monitors and lights were turned off in the recording room and adjacent control room to ensure that the environment was as dark as experimentally feasible. Luminance inside the recording chamber was less than 9x10^-4^ cd/m^2^ (SpectraScan PR 701S, Photo Research). Eye tracker and electrophysiological data were recorded while the animals executed eye-movements freely in the dark.

### Data Analysis

#### Saccade Identification

To identify saccade onset and offset times from the raw eye-tracker data, we used the Cluster Fix algorithm^94^. Cluster Fix performs k-means clustering on four parameters (distance, velocity, acceleration, and angular velocity) to find natural partitions in the eye-tracker data and identify saccades. To ensure that our set of identified eye movements was ‘clean,’ we imposed several additional criteria: 1) to avoid adjacent saccades affecting our analysis, we only considered saccades separated from neighboring saccades by at least 500 ms, 2) to exclude microsaccades, we only considered saccades that were greater in amplitude than 0.7 degrees of visual angle, 3) to exclude slower eye movements that Cluster Fix had mislabeled as saccades, we only considered eye movements less than 200 ms in duration, and 4) to exclude periods of data erroneously labeled as saccades due to blinking or drowsiness, we identified those periods using pupil diameter measurements and excluded nearby saccades.

Saccadic eye movements are known to have an approximately linear relationship between amplitude and peak velocity that is referred to as the main sequence^38^. To ensure that our approach identified a set of eye movements that quantitatively resembled the properties of saccades, we calculated the amplitude and peak velocity for each of the identified eye movements and found an approximately linear relationship (Figure 1A).

### Firing Rate Estimation and Modulation Index

To estimate the peri-saccadic firing rate, spikes were extracted for each saccade beginning 500 ms prior to saccade onset and ending 500 ms after saccade onset. To obtain a continuous estimate of the firing rate, each spike was convolved with a gaussian kernel and the average firing rate across saccades was calculated for each unit (Figure S1B; Figure S2). The kernel bandwidth was selected on a unit-by-unit basis at each point in time using a bandwidth optimization algorithm that minimized the mean integrated squared error^45^. In order to obtain reliable firing rate estimates, only units that fired at least 75 spikes within 500 ms of saccade onset (before or after) were included.

A modulation index was calculated for each unit to estimate the degree to which their activity deviated from baseline following saccade onset (Figure 2D, E). The modulation index was defined as *FR*_%_⁄*FR*_*b*_ − 1, where *FR*_*s*_ is the maximum firing rate in the 200 ms after saccade onset and *FR*_*b*_ is the baseline firing rate of the unit, calculated by averaging the firing rate for each unit in a time window 350 to 250 ms before saccade onset.

### Spike Count Correlations

To obtain a continuous estimate of spike count correlations, we calculated the Pearson correlation of spike counts across saccades at each time bin for every pair of simultaneously recorded single- and multi-units (Figure 4A, Figure S3A). The window was 201 ms centered in time and was shifted by 1 ms steps. Spike counts were z-scored to control for firing rate differences across time. Traces were smoothed with a 101 ms Gaussian weighted moving average. To perform the timing analysis, we took the local minimum in correlation as the time at which the rise begins. Bootstrapping was performed for each pair type to obtain confidence intervals on the estimate (Figure 4B).

### Spike-Spike Coherence

We computed the coherence between simultaneously recorded, multi-unit pairs within a cortical layer using multi-taper methods^95^ for two different windows: the 200 ms before and after saccade onset (Figure 4C). Spike trains were tapered with a single Slepian taper (TW = 3, K = 5). Magnitude of coherence estimates depend on the number of spikes used to create the estimates^96^. To control for differences in firing rate across the two windows, we adopted a rate-matching procedure^53^. In order to obtain a baseline for the coherence expected solely due to trends in firing time-locked to saccade onset, we also computed coherence in which saccade identities were randomly shuffled. Only units that elicited at least 50 spikes from each neuron cumulatively across all saccades in both time windows were considered. A modulation index, defined as (*SSC*_*after*_ − *SSC*_*before*_)⁄(*SSC*_*after*_ + *SSC*_*before*_), was calculated for each pair of units at each frequency band (Figure 4D).

### Power

Local field potential power was calculated with multi-tapered methods^95^. Signals were tapered with a single Slepian taper (TW = 3, K = 5). Spectrograms (Figure 5A) were generated by sliding a 201 ms window by increments of 1 ms. Power spectra before (-200 to 0 ms) and after (0 to 200 ms) saccade onset were calculated with static variants of the same methods (Figure 5B). No layer-specific differences were found, so results shown here are averaged across all recording channels.

### Pairwise Phase Consistency

The pairwise phase consistency (PPC) is an estimate of spike-field locking that is not biased by spike count or firing rate^97^. Two segments of data were analyzed: the 200 ms before and after saccade onset. In order to obtain reliable estimates, only units that fired at least 50 spikes cumulatively across all saccades during both time windows were considered. The spectro-temporal representation of the local field potential signal was generated through a continuous wavelet transform with a family of complex Morlet wavelets (spanning frequencies from 2 to 100 Hz, with 6 cycles at each frequency). Phase information was extracted at the time of each spike, and the PPC was then calculated for each unit with each LFP channel in the laminar probes (Figure 5C). We observed minimal change in the PPC as a function of LFP layer, so the results presented here were averaged across all recording channels.

### Computational Model

The firing rate or mean field model consists of a network of six populations representing excitatory (E) and inhibitory (I) populations in the superficial (L1), input (L2), and deep (L3) layers of the cortex. The firing rate model describes temporal changes in the firing rates (*r*) of each population as a function of the activity of two external inputs (*i*_*sacc*_, the saccadic signal, and *i*_*stim*_, the visual signal) and the activity of other populations in the network. The connectivity between populations is depicted in Figure 6A. The network consists of a coupled E-I network in each layer and captures the core aspect of connectivity between cortical layers: information flow from input to superficial to deep layer. The saccadic signal is sent to both E and I populations in the input layer, while the visual signal is sent only to the E population in the input layer. The saccadic signal is described by a ramping function (Figure 6C, top). The visual signal is only activated in the second set of simulations, where it is held constant to replicate sustained visual input. The six-population rate model is described by the following equations:

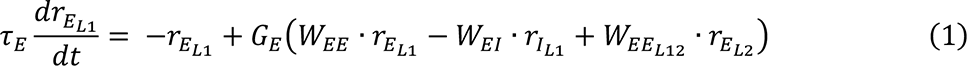

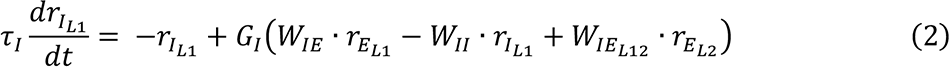

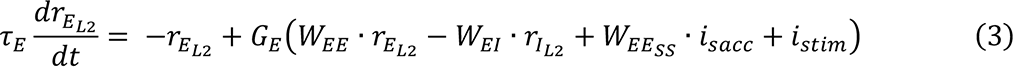

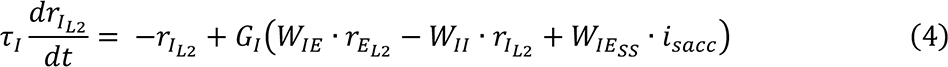

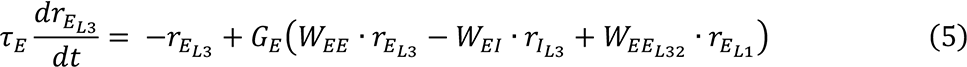

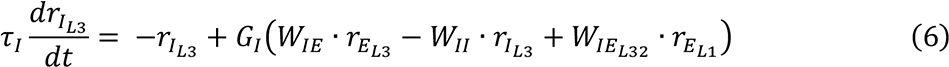

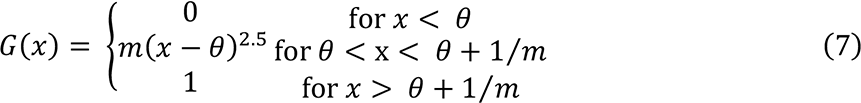

τ_*E*_ and τ_1_ are the rates at which the excitatory and inhibitory populations approach their steady states. *G*_*E*_ and *G*_1_ are the population response functions, described by equation (7) for the excitatory and inhibitory populations that transform the given inputs into a firing response. θ is the threshold input and *m* is the rate of the response function. Parameters tuned in the model to recapitulate the population level observations in experimental data: 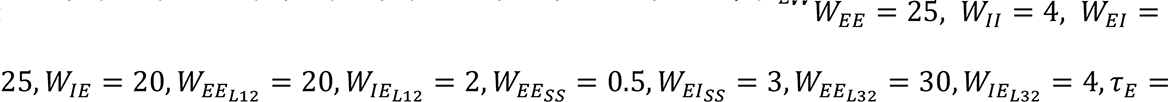 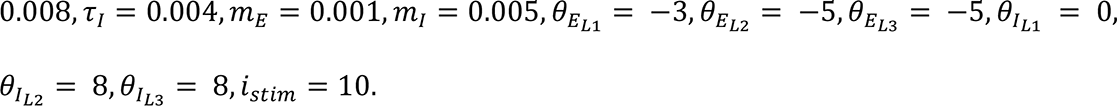

A further tuning was applied to the model to switch each E-I population between an inhibitory stabilized network (ISN) regime and a non-ISN one^60^. Experimental data from visual cortex suggest that a cortical network switches to an ISN regime during stimulus processing^59^, hence we included this feature to generate model predictions for saccadic suppression-related laminar activity during stimulus presentation. Previous modeling work has shown that a firing- rate E-I network model can switch between these regimes either by modification of connection weights (*W*), change in slope or rate of response functions (*m*) or a combination of both^60^. We implemented the simplest mechanism for the network to switch between ISN and non-ISN regime: nonlinear response functions. When visual stimulation (modeled as an excitatory input to the input layer E population) is applied to the model cortical column, the network moves to a higher slope region of its response functions, and switches to the ISN operating regime. A phase-plane analysis of the E-I network model illustrates how this results in a distinct response of the network to exactly the same saccadic signal input (Figure S4).

---

**Figure S1.**
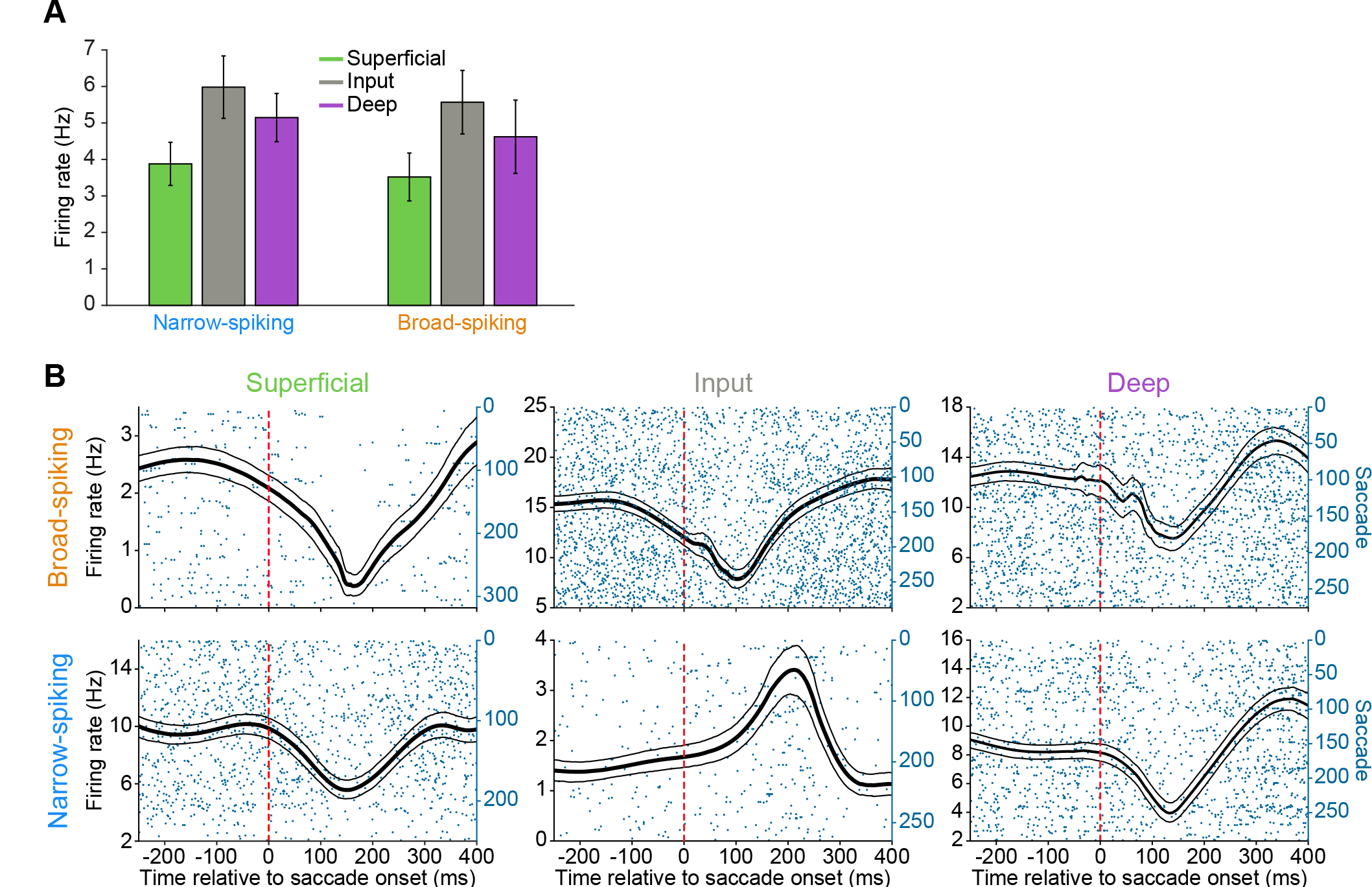
Peri-Saccadic Single-Unit Firing Rate Estimation. (A) Spontaneous firing rates of each neural subpopulation. (B) Peri-saccadic single-unit firing rates were estimated by smoothing individual spikes with gaussian kernels. The kernel bandwidth was selected for each unit at each point in time using an optimization algorithm that minimizes the mean integrated squared error^45^. The firing rate estimates produced by this approach are shown for six representative example units, one from each recorded neural subpopulation. The smoothed firing rates (black) are overlaid on the raster plots showing raw spikes (blue). In the raster plot, each row represents a single saccade. Thin lines are bootstrapped 95% confidence intervals of the firing rate estimate.

**Figure S2.**
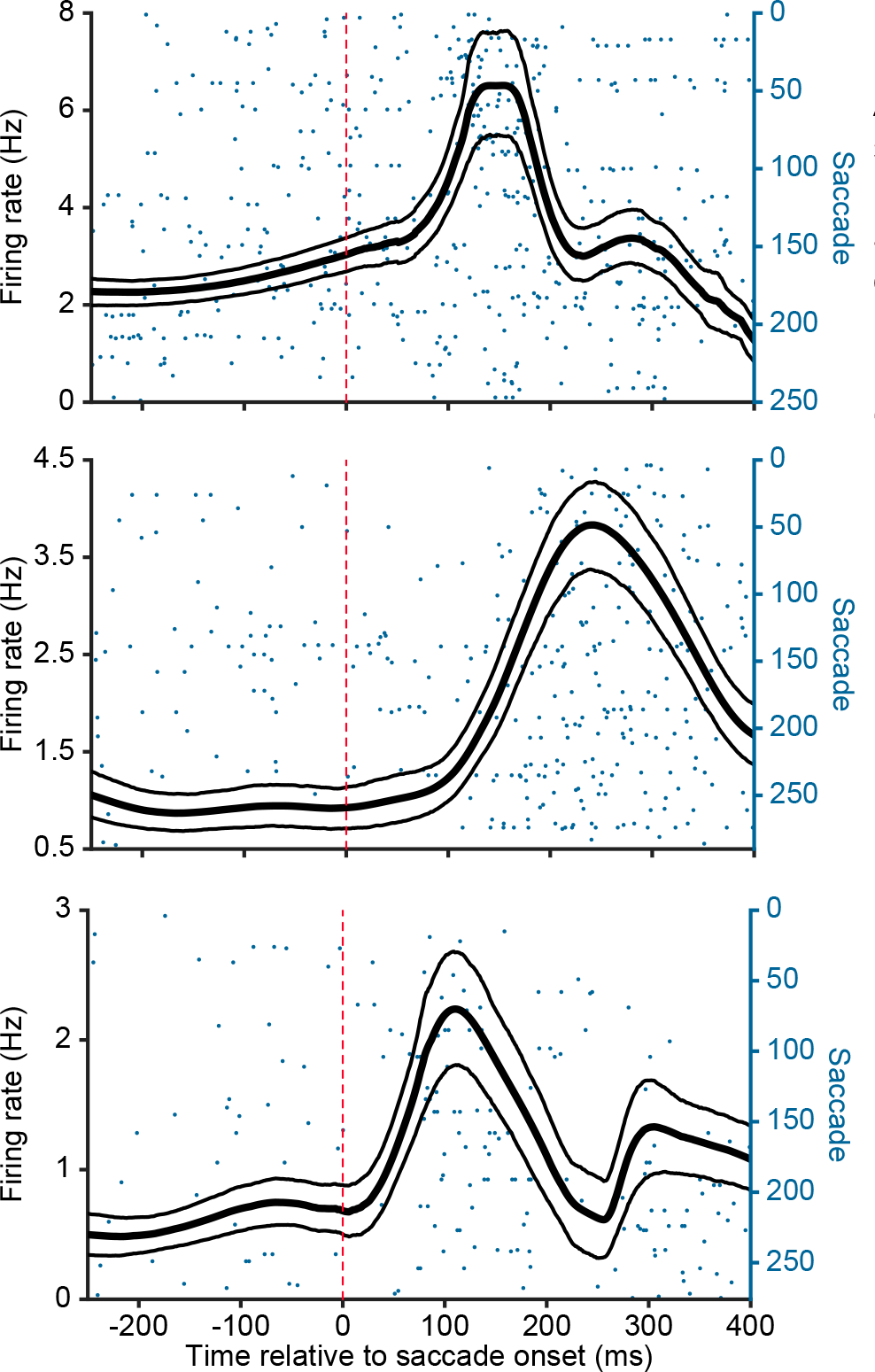
Additional Narrow-Spiking Input Layer Single-Unit Examples. Additional example narrow-spiking input layer single-units that display peri-saccadic enhancement of firing. Firing rates were estimated using a kernel smoothing approach and bandwidth optimization algorithm^45^. The smoothed firing rates (black) are overlaid on the spike raster plots (blue). Thin lines are bootstrapped 95% confidence intervals of the firing rate estimate.

**Figure S3.**
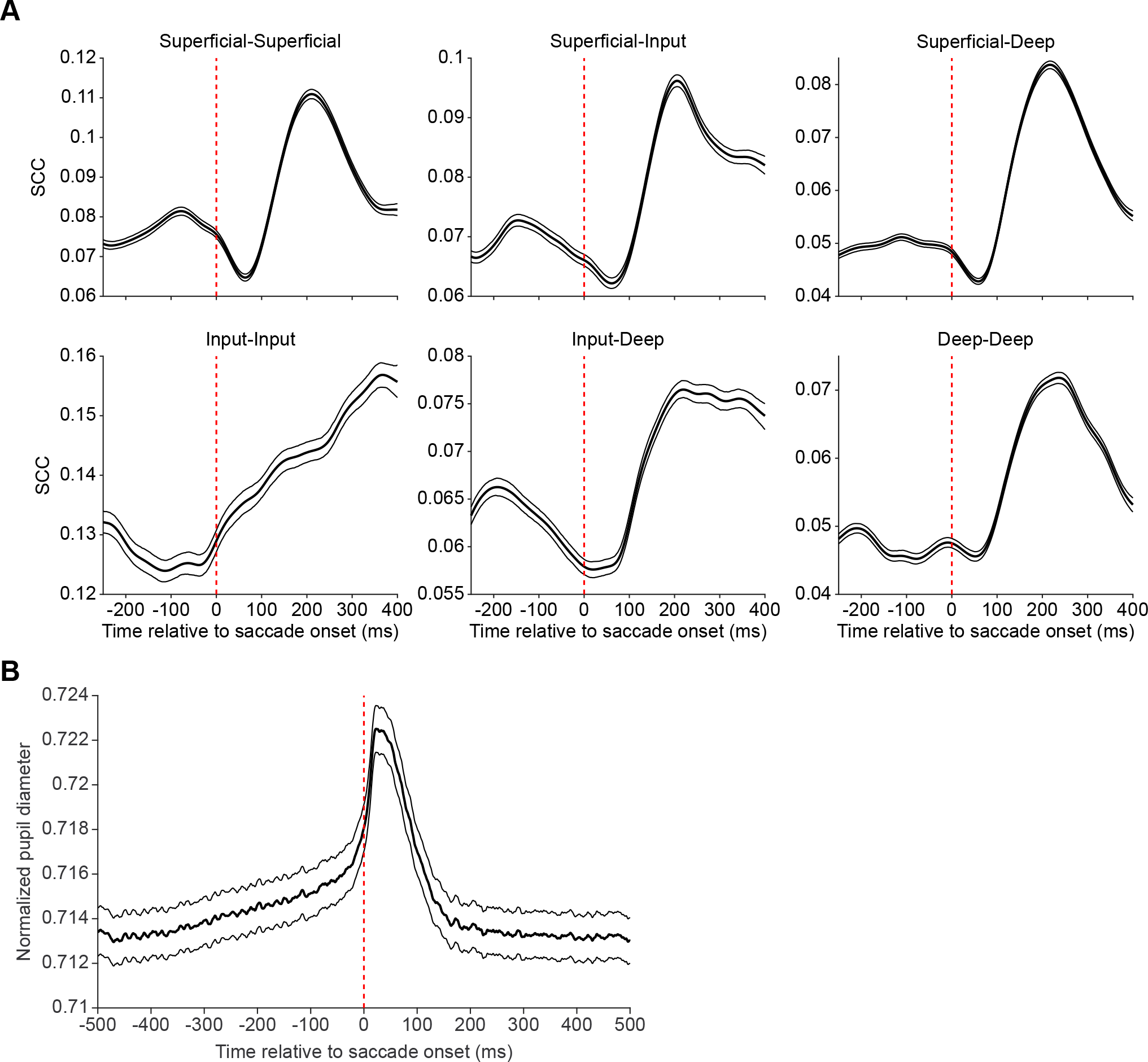
Input Layer Pairs Show Increases in Spike Count Correlations Prior to Saccade Onset. (A) Spike count correlations as a function of time relative to saccade onset for all combinations of layer-wise unit pairs. Superficial-superficial, *n* = 442 pairs; superficial-input, *n* = 558 pairs; superficial-deep, *n* = 755 pairs; input-input, *n* = 240 pairs; input-deep, *n* = 501 pairs; deep-deep, *n* = 680 pairs. Correlations were calculated with a sliding 201 ms window. Thin lines indicate bootstrapped 95% confidence intervals. (B) Normalized pupil diameter around the time of saccade onset (*n* = 4420 saccades). Thin lines indicate standard error of the mean.

**Figure S4.**
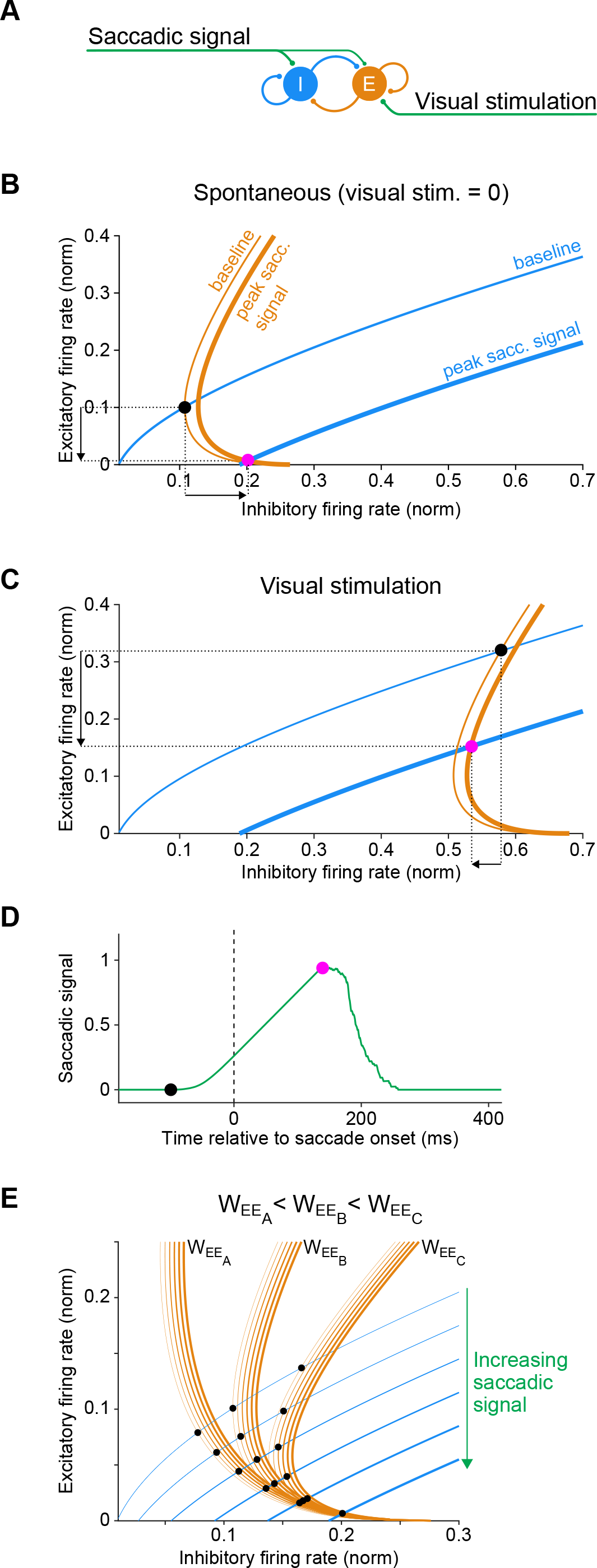
Phase Plane Analysis of the Input Layer E-I System. (A) The E-I network in the input layer. (B) The nullclines and stable activity levels of the input layer E and I populations during simulation of spontaneous activity. Equations and parameters are the same as those described in the Methods and used in Figure 6. The orange and blue curves show the E and I nullclines, respectively, of the E-I network in (A). Each point on a nullcline indicates the steady state firing rate of the E or I population when the activity of the other population is fixed, i.e. the rate of change in equations (3) or (4) is zero. The ISN and non-ISN regimes of the network are demarcated by the switch between positive and negative slopes along the E nullclines. This ability to switch is a consequence of strong E-E connectivity, and is absent when the E-E connectivity strength is either weak or zero. The thin nullclines depict firing rates when saccadic input is at baseline; the thick nullclines depict firing rates when saccadic input increases by a non-zero value. The intersections of the E and I nullclines indicate the steady state firing rates at which the network stabilizes if allowed to seek equilibrium (indicated by black and pink dots at baseline and during peak saccadic suppression, respectively). As illustrated here, the nullclines for our model intersect in the non-ISN regime in the absence of visual input, causing E and I activity to shift in opposite directions in response to the saccadic signal. (C) Same as in (B), but during simulated visual stimulation. The addition of excitatory input to the E population shifts the E nullclines to the right, causing them to intersect with the I nullclines in the ISN regime. As illustrated here, this causes the E and I populations to shift in the same direction in response to the saccadic signal. (D) The approximate level of saccadic signal strength that corresponds to the E (orange) and I (blue) nullcline sets shown in (B) and (C). The phase plane analysis is most applicable to constant levels of saccadic signal, and is only an approximate depiction of the scenario of a ramp signal shown in (D), and as used in the model simulation in Figure 6. (E) Effect of E-E connection strength on inhibitory firing rate changes in response to the saccadic signal. Each set of E nullclines corresponds to a given E-E connection strength (W_EE_), and illustrates the effects of raising saccadic signal magnitude (indicated by greater line thickness), causing either increases or decreases in steady state firing rates (black dots), as determined by their intersection with the I nullclines. This phase plane analysis approximately predicts three ways in which I firing rate can shift in response to increasing saccadic input: 1) it increases quickly in response to a small increase in saccadic signal (W_EEA_), 2) it increases minimally until a threshold level of saccadic signal is reached (W_EEB_), or 3) it first decreases then increases with rising saccadic signal strength (W_EEC_). The operating regime shown in the simulation results (Figure 6) that recapitulate experimental observations corresponds to the E-E connection strength illustrated by W_EEB_. In this case, the network is maintained at the boundary between ISN (positive E nullcline slope) and non-ISN (negative E nullcline slope) regimes in the absence of the saccadic signal.

## REFERENCES

1. Matin, E. Saccadic suppression: a review and an analysis. Psychol Bull 81, 899–917, doi:10.1037/h0037368 (1974).

2. Zuber, B. L. & Stark, L. Saccadic suppression: elevation of visual threshold associated with saccadic eye movements. Exp Neurol 16, 65–79, doi:10.1016/0014-4886(66)90087-2 (1966).

3. Thiele, A., Henning, P., Kubischik, M. & Hoffmann, K. P. Neural mechanisms of saccadic suppression. Science 295, 2460–2462, doi:10.1126/science.1068788 (2002).

4. Kleiser, R., Seitz, R. J. & Krekelberg, B. Neural correlates of saccadic suppression in humans. Curr Biol 14, 386–390, doi:10.1016/j.cub.2004.02.036 (2004).

5. Bremmer, F., Kubischik, M., Hoffmann, K. P. & Krekelberg, B. Neural dynamics of saccadic suppression. J Neurosci 29, 12374–12383, doi:10.1523/JNEUROSCI.2908-09.2009 (2009).

6. Zanos, T. P., Mineault, P. J., Guitton, D. & Pack, C. C. Mechanisms of Saccadic Suppression in Primate Cortical Area V4. J Neurosci 36, 9227–9239, doi:10.1523/JNEUROSCI.1015-16.2016 (2016).

7. Wurtz, R. H., Joiner, W. M. & Berman, R. A. Neuronal mechanisms for visual stability: progress and problems. Philos Trans R Soc Lond B Biol Sci 366, 492–503, doi:10.1098/rstb.2010.0186 (2011).

8. Sommer, M. A. & Wurtz, R. H. A pathway in primate brain for internal monitoring of movements. Science 296, 1480–1482, doi:10.1126/science.1069590 (2002).

9. Sommer, M. A. & Wurtz, R. H. Influence of the thalamus on spatial visual processing in frontal cortex. Nature 444, 374–377, doi:10.1038/nature05279 (2006).

10. Roe, A. W. et al. Toward a unified theory of visual area V4. Neuron 74, 12–29, doi:10.1016/j.neuron.2012.03.011 (2012).

11. Han, X., Xian, S. X. & Moore, T. Dynamic sensitivity of area V4 neurons during saccade preparation. Proc Natl Acad Sci U S A 106, 13046–13051, doi:10.1073/pnas.0902412106 (2009).

12. Hirsch, J. A. & Martinez, L. M. Laminar processing in the visual cortical column. Curr Opin Neurobiol 16, 377–384, doi:10.1016/j.conb.2006.06.014 (2006).

13. Douglas, R. J. & Martin, K. A. Neuronal circuits of the neocortex. Annu Rev Neurosci 27, 419–451, doi:10.1146/annurev.neuro.27.070203.144152 (2004).

14. Felleman, D. J. & Van Essen, D. C. in *Cereb cortex.* (Citeseer).

15. Ungerleider, L. G., Galkin, T. W., Desimone, R. & Gattass, R. Cortical connections of area V4 in the macaque. Cereb Cortex 18, 477–499, doi:10.1093/cercor/bhm061 (2008).

16. Rockland, K. S. & Pandya, D. N. Laminar origins and terminations of cortical connections of the occipital lobe in the rhesus monkey. Brain Res 179, 3–20, doi:10.1016/0006-8993(79)90485-2 (1979).

17. Boussaoud, D., Ungerleider, L. G. & Desimone, R. Pathways for motion analysis: cortical connections of the medial superior temporal and fundus of the superior temporal visual areas in the macaque. J Comp Neurol 296, 462–495, doi:10.1002/cne.902960311 (1990).

18. Distler, C., Boussaoud, D., Desimone, R. & Ungerleider, L. G. Cortical connections of inferior temporal area TEO in macaque monkeys. J Comp Neurol 334, 125–150, doi:10.1002/cne.903340111 (1993).

19. Borra, E., Ichinohe, N., Sato, T., Tanifuji, M. & Rockland, K. S. Cortical connections to area TE in monkey: hybrid modular and distributed organization. Cereb Cortex 20, 257–270, doi:10.1093/cercor/bhp096 (2010).

20. Gattass, R., Galkin, T. W., Desimone, R. & Ungerleider, L. G. Subcortical connections of area V4 in the macaque. J Comp Neurol 522, 1941–1965, doi:10.1002/cne.23513 (2014).

21. Anderson, J. C., Kennedy, H. & Martin, K. A. Pathways of attention: synaptic relationships of frontal eye field to V4, lateral intraparietal cortex, and area 46 in macaque monkey. J Neurosci 31, 10872–10881, doi:10.1523/JNEUROSCI.0622-11.2011 (2011).

22. Stanton, G. B., Bruce, C. J. & Goldberg, M. E. Topography of projections to posterior cortical areas from the macaque frontal eye fields. J Comp Neurol 353, 291–305, doi:10.1002/cne.903530210 (1995).

23. Benevento, L. A. & Rezak, M. The cortical projections of the inferior pulvinar and adjacent lateral pulvinar in the rhesus monkey (Macaca mulatta): an autoradiographic study. Brain Res 108, 1–24, doi:10.1016/0006-8993(76)90160-8 (1976).

24. Shipp, S. The functional logic of cortico-pulvinar connections. Philos Trans R Soc Lond B Biol Sci 358, 1605–1624, doi:10.1098/rstb.2002.1213 (2003).

25. Rockland, K. S. Distinctive Spatial and Laminar Organization of Single Axons from Lateral Pulvinar in the Macaque. Vision 4, 1 (2020).

26. McFarland, J. M., Bondy, A. G., Saunders, R. C., Cumming, B. G. & Butts, D. A. Saccadic modulation of stimulus processing in primary visual cortex. Nat Commun 6, 8110, doi:10.1038/ncomms9110 (2015).

27. Wurtz, R. H. Response of striate cortex neurons to stimuli during rapid eye movements in the monkey. J Neurophysiol 32, 975–986, doi:10.1152/jn.1969.32.6.975 (1969).

28. Bizzi, E. Discharge of frontal eye field neurons during saccadic and following eye movements in unanesthetized monkeys. Exp Brain Res 6, 69–80, doi:10.1007/BF00235447 (1968).

29. Schall, J. D. & Hanes, D. P. Neural basis of saccade target selection in frontal eye field during visual search. Nature 366, 467–469, doi:10.1038/366467a0 (1993).

30. Everling, S. & Munoz, D. P. Neuronal correlates for preparatory set associated with pro- saccades and anti-saccades in the primate frontal eye field. J Neurosci 20, 387–400 (2000).

31. Robinson, D. L., McClurkin, J. W., Kertzman, C. & Petersen, S. E. Visual responses of pulvinar and collicular neurons during eye movements of awake, trained macaques. J Neurophysiol 66, 485–496, doi:10.1152/jn.1991.66.2.485 (1991).

32. Berman, R. A. & Wurtz, R. H. Signals conveyed in the pulvinar pathway from superior colliculus to cortical area MT. J Neurosci 31, 373–384, doi:10.1523/JNEUROSCI.4738-10.2011 (2011).

33. Sudkamp, S. & Schmidt, M. Response characteristics of neurons in the pulvinar of awake cats to saccades and to visual stimulation. Exp Brain Res 133, 209–218, doi:10.1007/s002210000374 (2000).

34. Katzner, S., Busse, L. & Carandini, M. GABAA inhibition controls response gain in visual cortex. J Neurosci 31, 5931–5941, doi:10.1523/JNEUROSCI.5753-10.2011 (2011).

35. Wilson, N. R., Runyan, C. A., Wang, F. L. & Sur, M. Division and subtraction by distinct cortical inhibitory networks in vivo. Nature 488, 343–348, doi:10.1038/nature11347 (2012).

36. Mitchell, S. J. & Silver, R. A. Shunting inhibition modulates neuronal gain during synaptic excitation. Neuron 38, 433–445, doi:10.1016/s0896-6273(03)00200-9 (2003).

37. Berman, R. A., Cavanaugh, J., McAlonan, K. & Wurtz, R. H. A circuit for saccadic suppression in the primate brain. J Neurophysiol 117, 1720–1735, doi:10.1152/jn.00679.2016 (2017).

38. Bahill, A. T., Clark, M. R. & Stark, L. The main sequence, a tool for studying human eye movements. Mathematical Biosciences 24, 191–204 (1975).

39. Mitzdorf, U. Current source-density method and application in cat cerebral cortex: investigation of evoked potentials and EEG phenomena. Physiol Rev 65, 37–100, doi:10.1152/physrev.1985.65.1.37 (1985).

40. Nandy, A. S., Nassi, J. J. & Reynolds, J. H. Laminar Organization of Attentional Modulation in Macaque Visual Area V4. Neuron 93, 235–246, doi:10.1016/j.neuron.2016.11.029 (2017).

41. Mitchell, J. F., Sundberg, K. A. & Reynolds, J. H. Differential attention-dependent response modulation across cell classes in macaque visual area V4. Neuron 55, 131–141, doi:10.1016/j.neuron.2007.06.018 (2007).

42. Tamura, H., Kaneko, H., Kawasaki, K. & Fujita, I. Presumed inhibitory neurons in the macaque inferior temporal cortex: visual response properties and functional interactions with adjacent neurons. J Neurophysiol 91, 2782–2796, doi:10.1152/jn.01267.2003 (2004).

43. Hasenstaub, A. et al. Inhibitory postsynaptic potentials carry synchronized frequency information in active cortical networks. Neuron 47, 423–435, doi:10.1016/j.neuron.2005.06.016 (2005).

44. Kass, R. E., Ventura, V. & Cai, C. Statistical smoothing of neuronal data. Network 14, 5–15, doi:10.1088/0954-898x/14/1/301 (2003).

45. Shimazaki, H. & Shinomoto, S. Kernel bandwidth optimization in spike rate estimation. J Comput Neurosci 29, 171–182, doi:10.1007/s10827-009-0180-4 (2010).

46. Westheimer, G. Mechanism of saccadic eye movements. AMA Arch Ophthalmol 52, 710–724, doi:10.1001/archopht.1954.00920050716006 (1954).

47. Pare, M. & Munoz, D. P. Saccadic reaction time in the monkey: Advanced preparation of oculomotor programs is primarily responsible for express saccade occurrence. Journal of Neurophysiology 76, 3666–3681 (1996).

48. Deubel, H. & Schneider, W. X. Saccade target selection and object recognition: evidence for a common attentional mechanism. Vision Res 36, 1827–1837, doi:10.1016/0042-6989(95)00294-4 (1996).

49. Gilzenrat, M. S., Nieuwenhuis, S., Jepma, M. & Cohen, J. D. Pupil diameter tracks changes in control state predicted by the adaptive gain theory of locus coeruleus function. Cogn Affect Behav Neurosci 10, 252–269, doi:10.3758/CABN.10.2.252 (2010).

50. Reimer, J. et al. Pupil fluctuations track fast switching of cortical states during quiet wakefulness. Neuron 84, 355–362, doi:10.1016/j.neuron.2014.09.033 (2014).

51. Jainta, S., Vernet, M., Yang, Q. & Kapoula, Z. The pupil reflects motor preparation for saccades-even before the eye starts to move. Frontiers in human neuroscience 5, 97 (2011).

52. Doiron, B., Litwin-Kumar, A., Rosenbaum, R., Ocker, G. K. & Josic, K. The mechanics of state-dependent neural correlations. Nat Neurosci 19, 383–393, doi:10.1038/nn.4242 (2016).

53. Mitchell, J. F., Sundberg, K. A. & Reynolds, J. H. Spatial attention decorrelates intrinsic activity fluctuations in macaque area V4. Neuron 63, 879–888, doi:10.1016/j.neuron.2009.09.013 (2009).

54. Cohen, M. R. & Maunsell, J. H. Attention improves performance primarily by reducing interneuronal correlations. Nat Neurosci 12, 1594–1600, doi:10.1038/nn.2439 (2009).

55. Anastassiou, C. A., Perin, R., Markram, H. & Koch, C. Ephaptic coupling of cortical neurons. Nat Neurosci 14, 217–223, doi:10.1038/nn.2727 (2011).

56. Fries, P. Rhythms for Cognition: Communication through Coherence. Neuron 88, 220–235, doi:10.1016/j.neuron.2015.09.034 (2015).

57. Sirota, A. et al. Entrainment of neocortical neurons and gamma oscillations by the hippocampal theta rhythm. Neuron 60, 683–697, doi:10.1016/j.neuron.2008.09.014 (2008).

58. Veit, J., Hakim, R., Jadi, M. P., Sejnowski, T. J. & Adesnik, H. Cortical gamma band synchronization through somatostatin interneurons. Nature neuroscience 20, 951 (2017).

59. Ozeki, H., Finn, I. M., Schaffer, E. S., Miller, K. D. & Ferster, D. Inhibitory stabilization of the cortical network underlies visual surround suppression. Neuron 62, 578–592, doi:10.1016/j.neuron.2009.03.028 (2009).

60. Tsodyks, M. V., Skaggs, W. E., Sejnowski, T. J. & McNaughton, B. L. Paradoxical effects of external modulation of inhibitory interneurons. J Neurosci 17, 4382–4388 (1997).

61. Douglas, R. J. & Martin, K. A. Mapping the matrix: the ways of neocortex. Neuron 56, 226–238, doi:10.1016/j.neuron.2007.10.017 (2007).

62. Diamond, M. R., Ross, J. & Morrone, M. C. Extraretinal control of saccadic suppression. Journal of Neuroscience 20, 3449–3455 (2000).

63. Castet, E. & Masson, G. S. Motion perception during saccadic eye movements. Nature neuroscience 3, 177–183 (2000).

64. Campbell, F. W. & Wurtz, R. H. Saccadic omission: why we do not see a grey-out during a saccadic eye movement. Vision research 18, 1297–1303 (1978).

65. Burr, D. C., Morrone, M. C. & Ross, J. Selective suppression of the magnocellular visual pathway during saccadic eye movements. Nature 371, 511–513 (1994).

66. Bair, W., Zohary, E. & Newsome, W. T. Correlated firing in macaque visual area MT: time scales and relationship to behavior. J Neurosci 21, 1676–1697 (2001).

67. Kohn, A. & Smith, M. A. Stimulus dependence of neuronal correlation in primary visual cortex of the macaque. J Neurosci 25, 3661–3673, doi:10.1523/JNEUROSCI.5106-04.2005 (2005).

68. Shadlen, M. N. & Newsome, W. T. The variable discharge of cortical neurons: implications for connectivity, computation, and information coding. J Neurosci 18, 3870–3896 (1998).

69. Kriener, B., Tetzlaff, T., Aertsen, A., Diesmann, M. & Rotter, S. Correlations and population dynamics in cortical networks. Neural Comput 20, 2185–2226, doi:10.1162/neco.2008.02-07-474 (2008).

70. Stepniewska, I., QI, H.-X. & Kaas, J. H. Projections of the superior colliculus to subdivisions of the inferior pulvinar in New World and Old World monkeys. Visual neuroscience 17, 529–549 (2000).

71. Robinson, D. A. Eye movements evoked by collicular stimulation in the alert monkey. Vision research 12, 1795–1808 (1972).

72. Schiller, P. H., Sandell, J. H. & Maunsell, J. H. The effect of frontal eye field and superior colliculus lesions on saccadic latencies in the rhesus monkey. Journal of neurophysiology 57, 1033–1049 (1987).

73. Saalmann, Y. B., Pinsk, M. A., Wang, L., Li, X. & Kastner, S. The pulvinar regulates information transmission between cortical areas based on attention demands. Science 337, 753–756, doi:10.1126/science.1223082 (2012).

74. Zhou, H., Schafer, R. J. & Desimone, R. Pulvinar-Cortex Interactions in Vision and Attention. Neuron 89, 209–220, doi:10.1016/j.neuron.2015.11.034 (2016).

75. Petersen, S. E., Robinson, D. L. & Morris, J. D. Contributions of the pulvinar to visual spatial attention. Neuropsychologia 25, 97–105, doi:10.1016/0028-3932(87)90046-7 (1987).

76. Wilke, M., Turchi, J., Smith, K., Mishkin, M. & Leopold, D. A. Pulvinar inactivation disrupts selection of movement plans. J Neurosci 30, 8650–8659, doi:10.1523/JNEUROSCI.0953-10.2010 (2010).

77. Wilke, M., Kagan, I. & Andersen, R. A. Effects of pulvinar inactivation on spatial decision- making between equal and asymmetric reward options. J Cogn Neurosci 25, 1270–1283, doi:10.1162/jocn_a_00399 (2013).

78. Kawaguchi, Y. & Kubota, Y. Correlation of physiological subgroupings of nonpyramidal cells with parvalbumin- and calbindinD28k-immunoreactive neurons in layer V of rat frontal cortex. J Neurophysiol 70, 387–396, doi:10.1152/jn.1993.70.1.387 (1993).

79. Galarreta, M. & Hestrin, S. A network of fast-spiking cells in the neocortex connected by electrical synapses. Nature 402, 72–75, doi:10.1038/47029 (1999).

80. Cruikshank, S. J., Urabe, H., Nurmikko, A. V. & Connors, B. W. Pathway-specific feedforward circuits between thalamus and neocortex revealed by selective optical stimulation of axons. Neuron 65, 230–245, doi:10.1016/j.neuron.2009.12.025 (2010).

81. Gibson, J. R., Beierlein, M. & Connors, B. W. Two networks of electrically coupled inhibitory neurons in neocortex. Nature 402, 75–79, doi:10.1038/47035 (1999).

82. Agmon, A. & Connors, B. W. Correlation between Intrinsic Firing Patterns and Thalamocortical Synaptic Responses of Neurons in Mouse Barrel Cortex. Journal of Neuroscience 12, 319–329 (1992).

83. Poulet, J. F. & Petersen, C. C. Internal brain state regulates membrane potential synchrony in barrel cortex of behaving mice. Nature 454, 881–885, doi:10.1038/nature07150 (2008).

84. Vinck, M., Batista-Brito, R., Knoblich, U. & Cardin, J. A. Arousal and locomotion make distinct contributions to cortical activity patterns and visual encoding. Neuron 86, 740–754, doi:10.1016/j.neuron.2015.03.028 (2015).

85. Arroyo, S., Bennett, C. & Hestrin, S. Correlation of Synaptic Inputs in the Visual Cortex of Awake, Behaving Mice. Neuron 99, 1289–1301 e1282, doi:10.1016/j.neuron.2018.08.008 (2018).

86. John, E. R. et al. Invariant reversible QEEG effects of anesthetics. Conscious Cogn 10, 165–183, doi:10.1006/ccog.2001.0507 (2001).

87. Zohary, E., Shadlen, M. N. & Newsome, W. T. Correlated neuronal discharge rate and its implications for psychophysical performance. Nature 370, 140–143, doi:10.1038/370140a0 (1994).

88. Abbott, L. F. & Dayan, P. The effect of correlated variability on the accuracy of a population code. Neural Comput 11, 91–101, doi:10.1162/089976699300016827 (1999).

89. Bartolo, R., Saunders, R. C., Mitz, A. R. & Averbeck, B. B. Information-Limiting Correlations in Large Neural Populations. J Neurosci 40, 1668–1678, doi:10.1523/JNEUROSCI.2072-19.2019 (2020).

90. Averbeck, B. B., Latham, P. E. & Pouget, A. Neural correlations, population coding and computation. Nat Rev Neurosci 7, 358–366, doi:10.1038/nrn1888 (2006).

91. Lee, D. & Malpeli, J. G. Effects of saccades on the activity of neurons in the cat lateral geniculate nucleus. Journal of Neurophysiology 79, 922–936 (1998).

92. Nassi, J. J., Avery, M. C., Cetin, A. H., Roe, A. W. & Reynolds, J. H. Optogenetic Activation of Normalization in Alert Macaque Visual Cortex. Neuron 86, 1504–1517, doi:10.1016/j.neuron.2015.05.040 (2015).

93. Ruiz, O. et al. Optogenetics through windows on the brain in the nonhuman primate. J Neurophysiol 110, 1455–1467, doi:10.1152/jn.00153.2013 (2013).

94. Konig, S. D. & Buffalo, E. A. A nonparametric method for detecting fixations and saccades using cluster analysis: removing the need for arbitrary thresholds. J Neurosci Methods 227, 121–131, doi:10.1016/j.jneumeth.2014.01.032 (2014).

95. Mitra, P. P. & Pesaran, B. Analysis of dynamic brain imaging data. Biophys J 76, 691–708, doi:10.1016/S0006-3495(99)77236-X (1999).

96. Zeitler, M., Fries, P. & Gielen, S. Assessing neuronal coherence with single-unit, multi- unit, and local field potentials. Neural Comput 18, 2256–2281, doi:10.1162/neco.2006.18.9.2256 (2006).

97. Vinck, M., Battaglia, F. P., Womelsdorf, T. & Pennartz, C. Improved measures of phase- coupling between spikes and the Local Field Potential. J Comput Neurosci 33, 53–75, doi:10.1007/s10827-011-0374-4 (2012).

